# Aβ-induced distress of astrocytes triggers Alzheimer disease pathology through non-canonical δ secretase activity

**DOI:** 10.1101/2025.01.30.635618

**Authors:** Vanessa Schmidt, Ewelina Ziemlinska, Tomasz Obrebski, Ewa Zurawska-Plaksej, Jaroslaw Cendrowski, Barbara L Hempstead, Thomas E Willnow, Anna R Malik

## Abstract

The importance of astrocytes for Alzheimer disease (AD) pathology is increasingly appreciated, yet the mechanisms whereby this cell type impacts neurodegenerative processes remain elusive. In a genetic mouse model with diminished astrocyte stress response, even low levels of amyloid-β trigger astrocyte reactivity, resulting in brain inflammation and massive amyloid and tau pathologies. This dysfunctional response of astrocytes to amyloid-β acts through activation of δ secretase, a stress-induced protease implicated in both amyloid and tau-related proteolytic processing. Our findings identify a failed astrocyte stress response to amyloid-β as an early inducer of amyloid and tau co-morbidity, a noxious process in AD acting through a unique non-canonical secretase pathway.

## INTRODUCTION

Recent findings have identified the significance of glia for healthy brain aging, and disturbances in non-neuronal cell types as drivers of neurodegenerative processes in Alzheimer’s disease (AD). The underlying concepts have best been elucidated for microglia that play essential roles in phagocytic clearance of amyloid deposits as well as inflammatory processes in the AD brain (reviewed in (*1, 2*)). By contrast, the relevance of astrocytes, the major non-neuronal cell type in the brain for AD remains less well understood. Possible contributions to AD pathology are supported by recent evidence that astrocytes are sensitive to exposure to amyloid-β peptides (Aβ) (*3, 4*) and that astrocyte stress causes neuronal dysfunction and death (*5–7*). Also, the pro-inflammatory actions of reactive astrocytes in the AD brain likely contribute to progression of AD pathologies (*8–11*).

To elucidate novel molecular concepts of astrocytes (dys)function in AD, we focused on the sortilin-related receptor CNS expressed (SORCS) 2, a member of the VPS10P domain receptor family of intracellular sorting proteins (*12*). SORCS2 is a protective stress response factor in astrocytes, induced upon noxious insults such as ischemic stroke (*13*). Assuming a similarly protective function for SORCS2 in astrocyte stress response to Aβ, we investigated mouse models of AD lacking receptor expression. We hypothesized that lack of SORCS2 may sensitize astrocytes to amyloid burden and accentuate AD pathologies linked to astrocyte distress in the aging brain. In line with our hypothesis, SORCS2 deficiency aggravated αβ-induced stress and astrocyte cell death *in vitro* and *in vivo*, causing massive amyloid and tau pathologies in murine AD models. Surprisingly, astrocyte dysfunction was linked to aberrant activation of δ secretase, a stress-induced protease implicated in both amyloidogenic and tau-related proteolytic processing. Our findings identified astrocyte stress by Aβ as a causative mechanism of amyloid and tau comorbidities, noxious processes acting through a unique non-canonical secretase pathway.

## MATERIALS AND METHODS

### Mouse models

Mice with targeted disruption of *Sorcs2* (KO) have been described before (*14*). For analysis of human APP processing, wildtype (WT) and KO mice were crossed with the PDAPP line 109 (*15*). All animals were kept on an inbred C57BL/6N background and studied at 20 (young cohort) or 38-40 (old cohort) weeks of age. Astrocyte– and neuron-specific *Sorcs2* KO lines were generated by crossing the *Sorcs2^lox/lox^* line (*16*) with Cre transgenic strains Aldh1l1-Cre (JAX: #023748) or BAF53b-Cre (JAX: #027826). Subsequently, both lines were bred to PDAPP. Astrocyte-specific *Sorcs2* inactivation was induced in *PDAPP/Aldh1l1-Cre/Sorcs2^lox/lox^* animals by injection with 100 mg/kg body weight of tamoxifen (Sigma #T5648-1G) for five consecutive days at 10 – 12 weeks of age. All animal experimentations were performed according to institutional guidelines following approval by authorities of the State of Berlin (X9007/17; X9009/22; G0105/22) or the First Ethical Committee Warsaw (1375P1/2022).

### Analysis in mouse tissues

Expression analyses in mouse tissues by quantitative RT-PCR, Western blotting, ELISA, or immunohistology were performed using standard procedures as detailed in the supplementary methods. Cell death in brain tissues was determined as the amount of cytoplasmic histone-associated DNA fragments (mono– and oligonucleosomes) using the Cell Death Detection ELISA (Roche #11544675001) according to the manufactureŕs protocol. For δ secretase activity assays, brain tissues were homogenized in assay buffer (20 mM citric acid, 60 mM Na_2_HPO_4_, 1 mM DTT, 1 mM EDTA, 0.1% CHAPS, 0.5% Triton X-100, pH 4.5) and the protein concentrations adjusted to 50 μg/μl using the same buffer. Extracts (100 μl) were mixed with 100 μl of 20 μM δ secretase Z-AAN-AMC substrate (Bachem #4033201.0050) and the enzyme activity determined by fluorimetry (Ex: 346nm, Em: 443nm) over 2 hours.

### Flow cytometric analysis and quantification of brain cell types

Characterization of individual brain cell types was performed by FACS, following isolation of cells from adult mouse brains by Percoll gradients (see supplementary method for details).

Antibodies used for sorting were directed against markers of microglia and macrophages (CD45-BV421; 1:200, BD Biosciences #563890), endothelia (CD49a-FITC; 1:200, Miltenyi Biotec #130-107-636), oligodendrocytes (O4-PE; 1:200, Miltenyi Biotec #130-117-357) and astrocytes (ACSA2-APC, 1:100, Miltenyi Biotec #130-116-245; S100β-PE, 1:100, Novus Bio #NBP2-45267; GFAP-BV421, 1:100, Biolegend #644710; Aldh1l1-FITC, 1:100, Novus Bio #NBP2-50045F). Prior to staining for GFAP-BV421 and Aldh1l1-FITC, cells were fixed and permeabilized with Phosflow lyse/fix (BD Biosciences # 558049) and Perm Buffer III (BD Biosciences #558050) according to the manufacturers’ instructions.

### Assessing the response of primary astrocytes to Aβ

Astrocytes were isolated from newborn or adult mouse brain as described in supplementary methods. To determine Aβ uptake, primary astrocytes were treated with conditioned media from parental SY5Y cells or cell clone SY5Y-A, constitutively overexpressing human APP^695^ (*17*).

After 24 – 48 hours of incubation, levels of Aβ in astrocyte supernatants and lysates were measured by ELISA (Meso Scale Discovery). To determine viability, astrocytes were seeded in 8 technical replicates at a density of 5,000 cells per well in 96-well plates. Twenty-four hours after seeding, the medium was changed to conditioned media from cell lines SY5Y and SY5Y-A for 24 or 48 hrs. Finally, the medium was changed again to DMEM/HAMS F12 complete medium containing 1x Presto Blue reagent (Invitrogen #A13261). The fluorescence signal was measured after 60 min of incubation at excitation/emission wavelengths of 535/595 nm and ratio of (Aβ+) to (Aβ-) signals was calculated for each replicate.

For analysis of the lysosomal compartment, astrocytes were seeded at a density of 5,000 –7,000 cells per well in 96-well plates for 24 hrs. Then, the medium was switched to conditioned media from cell lines SY5Y and SY5Y-A for 24 hrs. Then, cells were live-stained for 20 min with 50 nM LysoTracker Red DND-99 (L7528; Thermo Fisher Scientific) and Hoechst and scanned using Opera Phenix high content screening microscope (PerkinElmer) with 40 × 1.1 NA water immersion objective. The Harmony 4.9 software (PerkinElmer) was used for image acquisition and quantitative analysis. More than 10 microscopic fields were analysed for each experimental condition to quantify lysosome number and cumulative intensity of their fluorescence. The average number of cells used for analysis was between 350 and 450 per condition. Maximum intensity projection images were obtained from three to five Z-stack planes with 1 μm interval. Images were assembled in ImageJ and Photoshop (Adobe) with only linear adjustments of contrast and brightness.

### Statistics

For all experiments, an indicated number n is the number of mice per group used in an experiment. Each mouse represents a statistically independent experimental unit, which was treated accordingly as an independent value in the statistical analysis. Statistical analyses were performed using GraphPad Prism 8 Software. Data are presented as mean ± standard error of the mean (SEM), mean ± standard deviation (SD) or geometric mean ± confidence interval (CI) as indicated in the respective figure legends. To compare two groups, unpaired two-sided Student’s *t* test or unpaired two-sided Mann–Whitney U test was used, depending on normal distribution of data. For experiments with more than two parameters, an ordinary or repeated-measures Two-way ANOVA with Tukeýs or Holm-Sidak’s multiple comparison test was applied. Before selecting a test, a preliminary analysis for normal distribution of the data was performed. The exact test used is indicated in the figure legends.

## RESULTS

### SORCS2 deficiency sensitizes astrocytes to amyloid-induced cell death

Initially, we tested our assumption that deficiency for the stress response factor SORCS2 may impact the ability of astrocytes to cope with noxious stimuli by Aβ peptides. Treatment of primary murine astrocytes with Aβ-conditioned medium from SY5Y cells, overexpressing a human *APP* (*17*), induced *Sorcs2* transcript levels, linking receptor expression with Aβ sensing (Fig. 1A). Exposure to Aβ-conditioned medium increased clearance (Fig. 1B) and intracellular accumulation of Aβ40 and Aβ42 in astrocytes from mice carrying a targeted disruption of *Sorcs2* (KO) (*14*) compared to wildtype (WT) cells, as shown by fluorescence microscopy (Fig. 1C) and ELISA (Fig. 1D). Intracellular accumulation of Aβ coincided with elevated levels of glial fibrillary acid protein (GFAP) (Fig. 1E) and a decrease in lysosomal acidification (Fig. 1F), stress responses more pronounced in KO than WT astrocytes. Ultimately, primary KO astrocytes reacted to prolonged Aβ exposure with decreased viability as shown by PrestoBlue assay (Fig. 1G). Reduced viability coincided with enhanced apoptosis as judged from increased levels of cleaved forms of poly ADP-ribose polymerase (PARP) and cleaved caspase 3 in KO as compared to WT astrocytes (Fig. 1H-I). Taken together, these data supported a presumed role for SORCS2 in mitigating noxious insults imposed on astrocytes by Aβ, and they suggested SORCS2 deficiency as useful experimental paradigm to investigate consequences of heightened astrocyte stress in AD.

**Figure 1:**
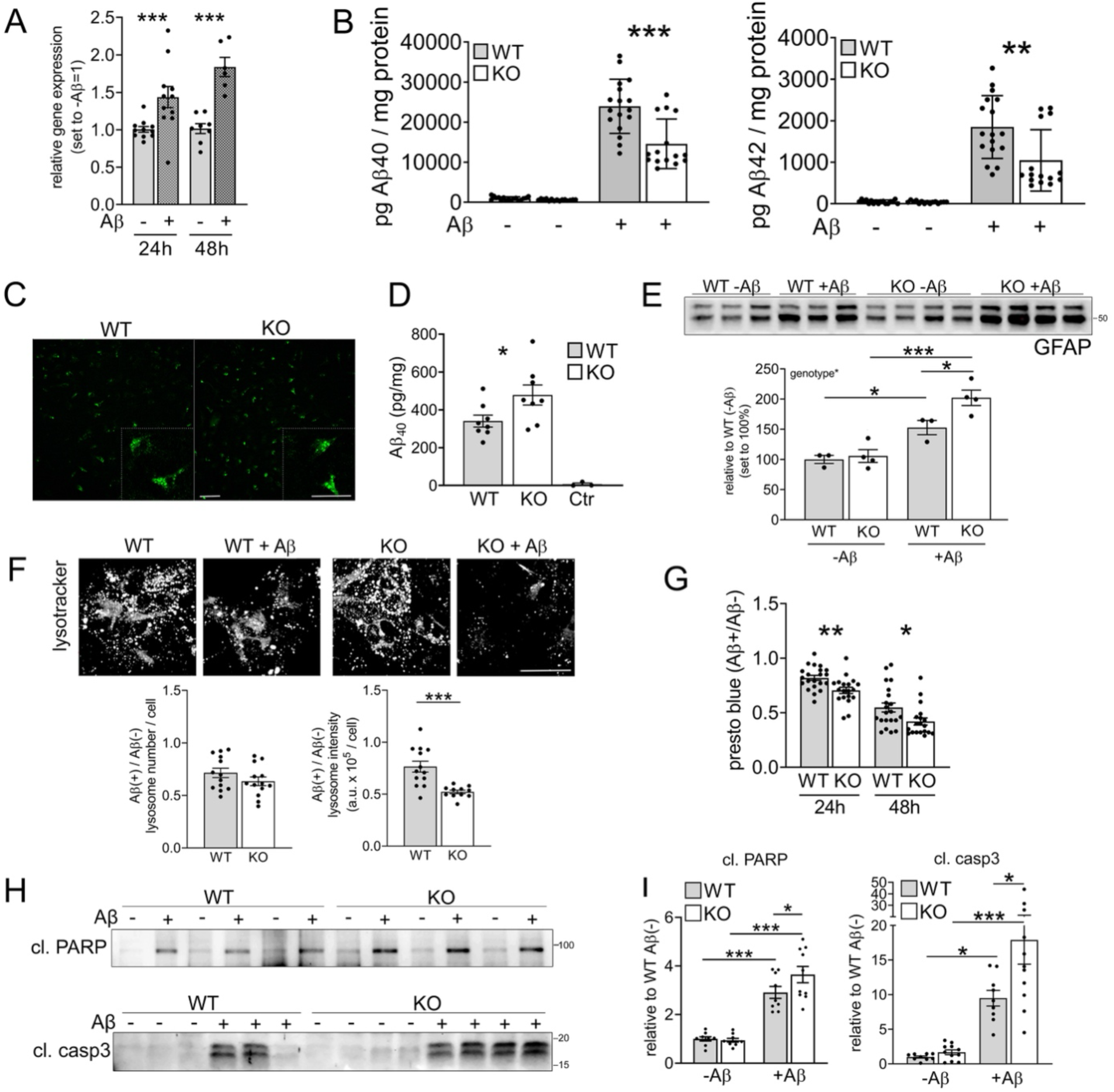
SORCS2-deficient primary astrocytes exhibit enhanced cell stress in response to Aβ (**A**) Levels of *Sorcs2* transcripts as determined by qRT-PCR in primary astrocytes from wildtype (WT) mice treated with control (−) or Aβ-conditioned media (+) for 24h or 48h. Aβ-conditioned medium had been generated from SY5Y cells stably overexpressing human APP (*17*). Data are expressed as relative expression levels (set to 1 for –Aβ) and given as mean ± SEM from n=6-11 independent cell preparations per group (unpaired Mann-Whitney U test). **(B)** Primary WT or SORCS2-deficient (KO) astrocytes were treated with control (−) or Aβ-conditioned medium (+) and residual levels of Aβ were measured after 24h using ELISA. Media from KO astrocytes showed lower levels of Aβ_40_ and Aβ_42_, indicating increased cellular Aβ uptake. Data are given as mean ± SEM from n=15-17 biological replicates (unpaired Mann-Whitney U-test). **(C, D)** Primary WT and KO astrocytes were treated with 1 µM HiLyte-Aβ_40_ for 6h. Aβ uptake was visualized by fluorescence microscopy for HiLyte-Aβ_40_ (C, green) and quantified by ELISA in cell lysates (D). In D, intracellular Aβ levels are higher in KO as compared to WT cells. No Aβ is detected in WT cells treated with control medium (Ctr). Data are given as mean ± SEM from n=8 animals per group (two-sided unpaired Student’s *t*-test). Scale bar: 50 μm (inset: 200 μm)**. (E)** Western blot analysis of glial fibrillary acidic protein (GFAP) levels in WT and KO astrocytes after 48h of treatment with control (−) or Aβ-conditioned (+) medium (upper panel) and quantification thereof (lower panel). Data are expressed as levels relative to WT –Aβ (set to 100%) and given as mean ± SEM from n=3-4 independent cell preparations per genotype and condition (ordinary two-way ANOVA with Tukeýs multiple comparisons test to test significance for genotype). **(F)** Primary WT or KO astrocytes were cultured for 24h in the presence or absence of Aβ-conditioned medium followed by live staining with LysoTracker Red dye (lysotracker). Lysotracker fluorescence signals (upper panel) were used to quantify lysosome numbers as well as cumulative signal intensity per cell (lower panel; in arbitrary units, a.u.). In KO, and to a lesser extent in WT cells, Aβ treatment decreases lysotracker intensities without impacting lysosome numbers, suggesting impaired lysosomal acidification. Data are given as mean ± SEM from n=13 independent cell preparations per genotype (two-sided unpaired Student’s *t*-test). Scale bar: 50 μm. **(G)** Cell viability was measured using presto blue in primary astrocytes treated with control or Aβ-containing medium for 24h and 48h. Data are given as mean ± SEM of fluorescent signal ratio (+Aβ)/(−Aβ) from n=19-21 independent cell preparations per genotype and condition (two-sided unpaired Student’s *t*-test). Aβ treatment decreased viability in KO to a higher extent than in WT cells. **(H-I)** Western blot analyses of cleaved poly (ADP-Ribose) polymerase 1 (cl. PARP) and cleaved caspase 3 (cl. casp3) levels in WT and KO astrocytes after 24h (cl. PARP) or 48h (cl. casp3) of treatment with control (−Aβ) or Aβ medium (+Aβ). Exemplary blots from 3 independent experiments per condition (H) as well as quantification from densitometric scanning of replicate blots (I) are given. Levels are expressed as relative to WT –Aβ (set to 1), and are given as mean ± SEM from n=9-12 independent cell preparations per genotype and condition (ordinary Two-way ANOVA with Holm-Sidak’s multiple comparisons test). *, p<0.05; **, P < 0.01; ***, P < 0.001

### Loss of SORCS2 results in amyloid and tau co-morbidity in a mouse model of AD

To interrogate the impact of enhanced astrocyte stress on AD pathology, we crossed *Sorcs2*^-/-^ mice with transgenic animals expressing the human *APP^Ind^* transgene under control of the platelet-derived growth factor β promoter (PDAPP strain), an established model of AD (*15*). The resulting (PDAPP x *Sorcs2*^-/-^) mice are referred to as PDAPP/KO herein. They were compared to sex– and age-matched (PDAPP x *Sorcs2*^+/+^) animals (PDAPP/WT).

PDAPP/KO females showed an age-dependent increase in brain cortex levels of soluble Aβ_40_ and Aβ_42_ compared to PDAPP/WT, starting from 12 weeks of age (Fig. 2A-B). At 20 weeks of age, cortical Aβ_40_ and Aβ_42_ levels displayed a bimodal distribution in the PDAPP/KO cohort with half of the animals having a 2-fold and the other half an approximately 9-fold increase over PDAPP/WT (Fig. 2C). At 40 weeks of age, all PDAPP/KO mice showed massive accumulation of Aβ_40_ and Aβ_42_ with cortex levels exceeding those in PDAPP/WT by more than 10-fold for Aβ_40_ and 30-fold for Aβ_42_ (Fig. 2D). The same bimodal distribution at 20 weeks, and massive accumulation in all mutant mice at 40 weeks of age, was observed for soluble Aβ_40_ and Aβ_42_ in hippocampal extracts (Fig. 2E-F). Aβ accumulation resulted in pronounced senile plaque deposition in cortex and hippocampus of aged PDAPP/KO females, not seen in controls (Fig. 2G).

**Figure 2:**
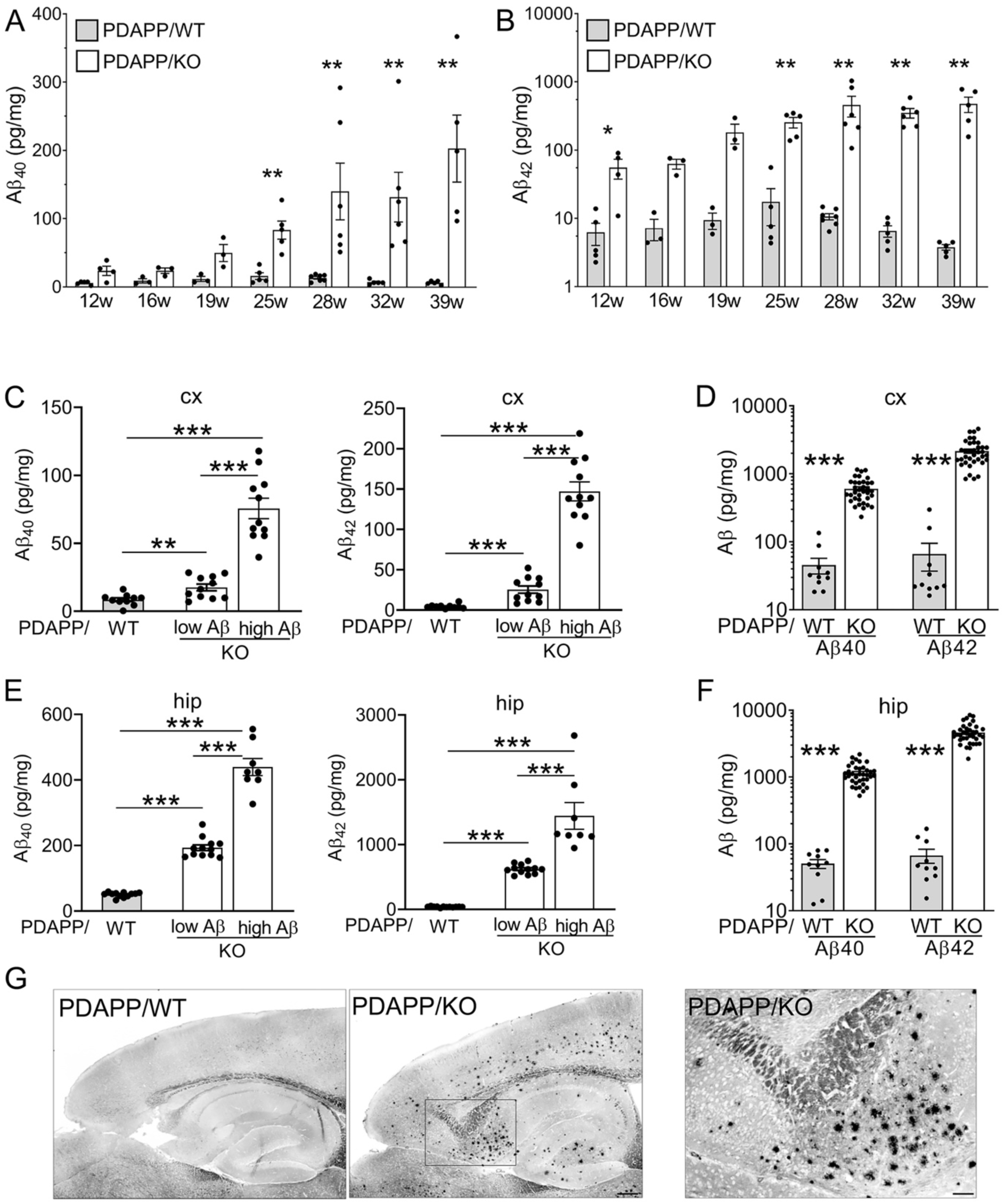
SORCS2 deficiency causes massive age-dependent brain amyloidosis in a mouse model of AD **(A, B)** Age-dependent increase in soluble Aβ_40_ (A) and Aβ_42_ (B) levels in cortex of PDAPP/KO females as compared to PDAPP/WT controls. Data are given as mean ± SEM from n=3-7 animals per genotype (unpaired Mann-Whitney U test). **(C, D)** Levels of soluble Aβ_40_ and Aβ_42_ in cortex (cx) of PDAPP/WT and PDAPP/KO mice at 20 (C) and 40 weeks of age (D). Massive increase in Aβ levels is seen for some PDAPP/KO animals at 20 weeks, but for all animals at 40 weeks of age. Data are given as mean ± SEM from n=10-36 animals per group (two-sided unpaired Student’s *t*-test for C, unpaired Mann-Whitney U test for D). **(E, F)** Experiment as in C and D but using hippocampal (hip) extracts of young (20 weeks, E) and aged (40 weeks, F) PDAPP/WT and PDAPP/KO females. Data are given as mean ± SEM from n=8-36 animals per group (unpaired two-sided Student’s *t*-test in E, unpaired Mann-Whitney U test in F). Note logarithmic scales in B, D, and F. *, P < 0.05; **, P < 0.01; ***, P < 0.001. **(G)** Thioflavin S-stained sagittal brain sections document amyloid plaques in hippocampal and cortical regions of 40 weeks old PDAPP/KO as compared to PDAPP/WT females. Representative sections from a total of 3 animals per genotype are shown. The square in the overview image indicates the area of the PDAPP/KO tissue magnified to the right. Scale bar: 200 μm (overview), 40 μm (zoom-in).

Remarkably, loss of SORCS2 also caused tauopathy-like phenotypes in PDAPP/KO females. In detail, SORCS2 deficiency increased levels of murine tau phosphorylated at inorganic pyrophosphatase 2 (PPA2) target sites Ser_202_ and Thr_205_ (antibody AT8), in cortex (Fig. 3A) and hippocampus (Fig. 3B) of aged PDAPP/KO mice, as shown by Western blot. Levels of murine ptau_Thr231_ (AT180 antibody) were also increased, while the levels of total tau (HT7 antibody) were unchanged compared to control females (Fig. 3A-B). A relative increase in the ratio of ptau_Thr231_ over total tau in cortex and hippocampus of PDAPP/KO brains was confirmed by ELISA (Fig. 3C-D). Both ptau_Ser202/Thr205_ and ptau_Thr231_ variants contribute to formation of tau tangles (*18–22*). Immunohistology confirmed elevated levels of ptau_Ser202/Thr205_ immunoreactivity in several brain regions of PDAPP/KO compared to PDAPP/WT females (Fig. 3E-F).

**Figure 3:**
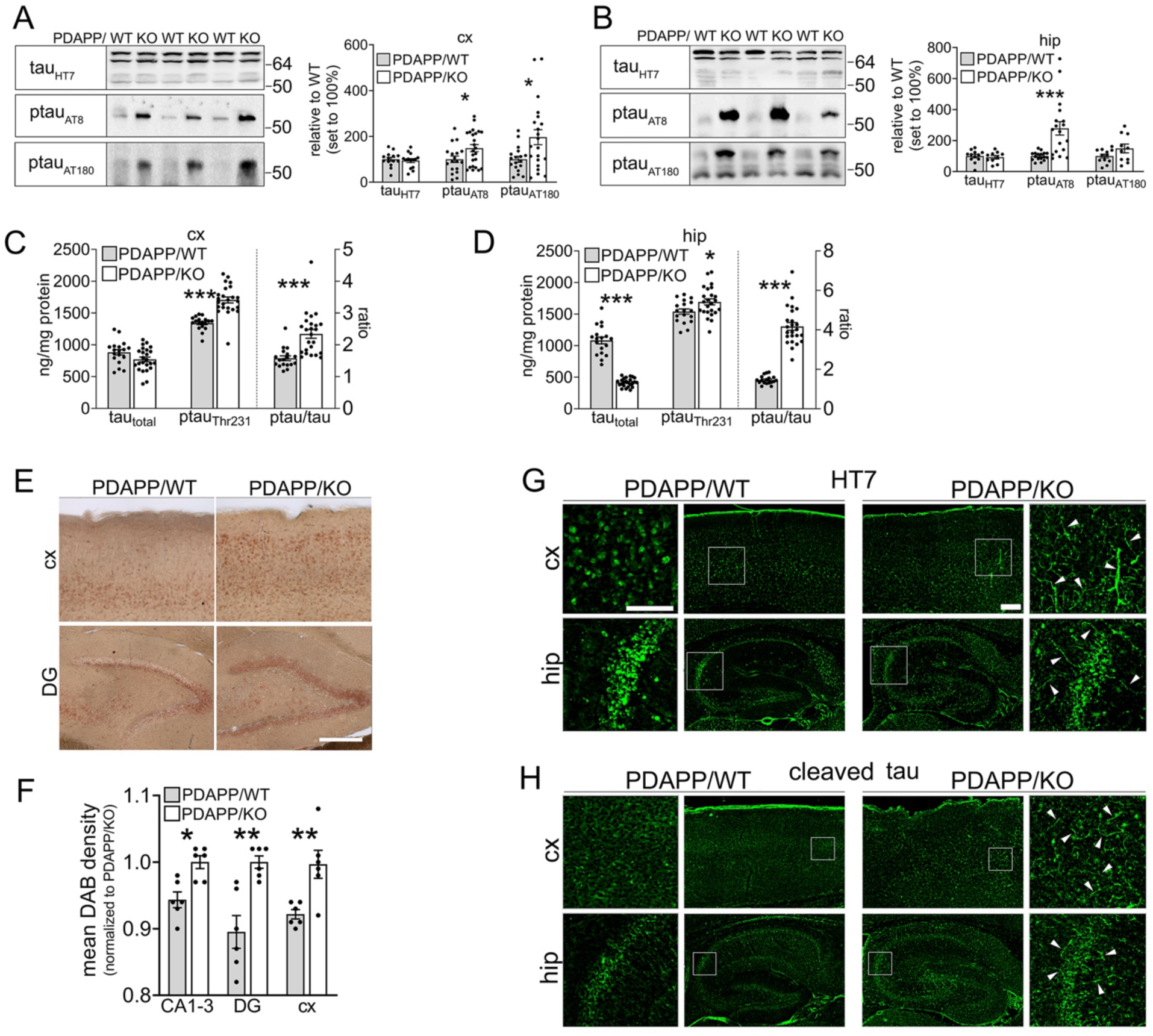
SORCS2 deficiency induces tau pathology in a mouse model of AD **(A, B)** Representative Western blot analyses, and densitometric quantification of replicate blots thereof, document levels of total tau (HT7) as well as phosphorylated variants ptau_Σερ202/Thr205_ (AT8) and ptau_Thr231_ (AT180) in cortical (A) and hippocampal (B) brain extracts of 40 weeks old PDAPP/WT or PDAPP/KO females. Data are expressed as relative to WT (set to 100%) and given as mean ± SEM from n=10-14 (HT7) and n= 10-24 (AT8 and AT180) animals per genotype (two-sided unpaired Student’s *t*-test). **(C, D)** Levels of total (tau_τοταλ_) and ptau_Thr231_, as well as ratio of ptau_Thr231_/tau_τοταλ,_ in cortex (C) and hippocampus (D) of 40 weeks old female mice as determined by ELISA. Data are given as mean ± SEM from n=18-24 animals per genotype (two-sided unpaired Student’s *t*-test). **(E, F)** Immunostaining for ptau _Σερ202/Thr205_ (AT8 antibody) on histological sections of cortical (cx) and hippocampal (hip) brain regions of 40 weeks old PDAPP/WT and PDAPP/KO female mice. Panel E depicts exemplary images of a total six animals per genotype. Panel F gives the quantification of ptau _Σερ202/Thr205_ levels based on mean 3,3’-diaminobenzidine (DAB) intensities in 5 sections each from 6 animals per genotype. Data are given as mean ± SEM (unpaired Mann-Whitney U-test). Scale bar, 200 μm. *, P < 0.05; **, P < 0.01; ***, P < 0.001 **(G, H)** Representative images of cx and hip sections from 40 weeks old female PDAPP/WT and PDAPP/KO mice immunostained for total tau (G, HT7 antibody) and cleaved forms of tau_ασπ421, Asp422_ (H, tauC3). Images are given as overview images and higher magnification zoom (indicated as white squares in the overviews). Arrowheads indicate fibrillary appearance of tau and tauC3 immunoreactivity in PDAPP/KO tissue. Scale bar: 150 μm (inset), 200 μm (overview).

Immunoreactivity for total tau (HT7) or cleaved tau (tau_Asp421/Asp422_), which rapidly assembles into filaments (*23, 24*), showed a distinct fibrillary appearance in PDAPP/KO brains, not seen in PDAPP/WT tissue (Fig. 3G-H). Levels of ptau_Ser202/Thr205_ and ptau_Thr231_ were unchanged in cortex (Fig. S1A-B) and hippocampus (Fig. S1C-D) of aged female KO mice lacking the PDAPP transgene. Also, the ratio of ptau_Thr231_ over total tau in cortex and hippocampus was comparable to that in WT brains (Fig. S1E-F). These findings documented that stress from human Aβ in the PDAPP line was necessary to trigger tau hyperphosphorylation in KO females. Massive accumulation of Aβ_40_ and Aβ_42_ in cortex and hippocampus was also seen in PDAPP/KO males at 40 weeks of age (Fig. S2A). However, contrary to females, no significant increases in ptau_Ser202/Thr205_ and ptau_Thr231_ levels (Fig. S2B-C) or change in ptau_Thr231_/tau ratio (Fig. S2D-E) were seen, in line with a bias for tau pathology in females (*25, 26*).

### Aβ stress induces astrocyte reactivity and loss in SORCS2-deficient mice

Next, we investigated the cellular consequences of amyloid and tau co-morbidity in our mouse models. Using a cell death assay, no difference in viability was seen comparing cortical or hippocampal extracts of WT and KO female brains lacking *APP*_Ind_ (Fig. 4A). Cell death increased in both genotypes in the presence of the PDAPP transgene, but this increase was significantly higher in PDAPP/KO than PDAPP/WT females (Fig. 4A), indicating heightened sensitivity of SORCS2-deficient brains to Aβ-induced cell death.

**Figure 4:**
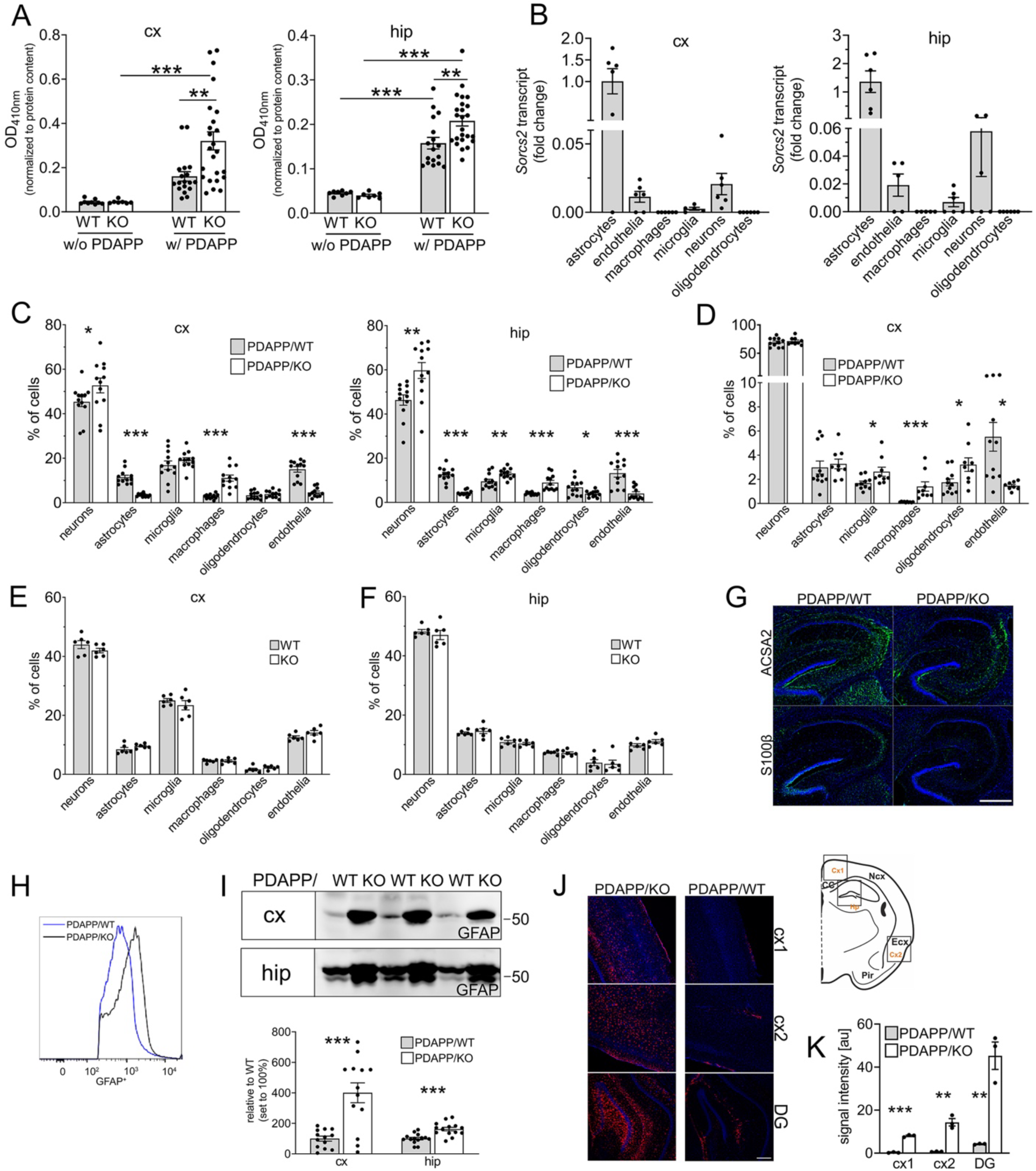
Aβ induces astrogliosis and loss of astrocytes in SORCS2-deficient mice (**A**) Nucleosome fragmentation assay documents increased cell death in cortical (cx, left panel) and hippocampal (hip, right panel) brain extracts of 40 weeks old PDAPP/WT and PDAPP/KO females when compared to WT and KO controls lacking PDAPP. However, PDAPP-induced cell death was significantly higher in PDAPP/KO than PDAPP/WT. Data are given as mean ± SEM from n=18-24 (PDAPP) and n=7-9 (non-PDAPP) animals per genotype group (ordinary two-way ANOVA with Tukeýs multiple comparisons test). (**B**) Transcript levels for *Sorcs2* in the indicated cell types isolated by FACS from cx (left panel) and hip (right panel) of PDAPP/WT females. Data are given as mean ± SEM from n=6 animals per genotype. **(C)** Cell type distribution in cx (upper panel) and hip (lower panel) of 40 weeks old PDAPP/WT and PDAPP/KO females as determined by FACS. PDAPP/KO animals show a significant reduction in relative numbers of astrocytes and endothelial cells, and a concomitant increase in microglia and macrophages. Data are given as mean ± SEM from n=11-12 animals per genotype (two-sided unpaired *t*-test). **(D)** Cell type distribution in cx of 20 weeks old PDAPP/WT and PDAPP/KO females based on FACS. Relative abundance of astrocytes is unchanged, while numbers of microglia and macrophages increase in young PDAPP/KO mice. Data are given as mean ± SEM from n=9-11 animals per genotype (unpaired Mann-Whitney U-test). **(E-F)** FACS documenting relative distribution of various cell types to be unchanged comparing cx (E) and hip (F) of 40 weeks old WT and KO females lacking PDAPP. Data are given as mean ± SEM from n=6 animals per genotype (unpaired Mann-Whitney U-test). **(G)** Immunostaining of hip regions of 40 weeks old PDAPP/WT and PDAPP/KO females for astrocyte markers ACSA2 and S100β (green). Nuclei were counterstained with DAPI (blue). Scale bar, 400 μm. **(H)** Representative FACS histogram of GFAP levels in sorted astrocytes documenting higher levels in aged PDAPP/KO as compared to PDAPP/WT female brains. (**I**) Representative Western blot for GFAP in brain extracts (upper panel), and quantification from densitometric scanning of replicate blots (lower panel), show increased levels in cx and hip of 40 weeks PDAPP/KO females compared to matched PDAPP/WT controls. Data are expressed as relative to WT (set to 100%) and given as mean ± SEM from n=13-14 animals per genotype (two-sided unpaired Student’s *t*-test). **(J)** Representative images of GFAP immunostaining (red) in brain regions of 40 weeks old PDAPP/KO as compared to PDAPP/WT female mice. A schematic depicting analyzed cortical (cx1, cx2) and dentate gyrus (DG) regions is shown. Nuclei were counterstained with DAPI (blue). Scale bar: 200 μm. (**K**) GFAP signal intensities (arbitrary units, au) quantified from replicate immunostainings exemplified in J, show significant increase in GFAP levels in indicated brain regions in PDAPPP/KO females compared to PDAPP/WT controls. Data are given as mean ± SEM from n=3 animals per genotype (two-sided unpaired Student’s *t*-test).*, P < 0.05; **, P < 0.01; ***, P < 0.001

To identify the cell type(s) most impacted by Aβ stress, we compared the cell composition of brains in aged PDAPP/WT and PDAPP/KO females. To do so, we established a FACS protocol to isolate pure populations of individual cell types, including neurons, astrocytes, microglia, and oligodendrocytes (Fig. S3). Quantitative RT-PCR on sorted cell populations documented *Sorcs2* transcripts to mainly localize to astrocytes, and to a much lesser extent to neurons and endothelial cells, in cortex and hippocampus of the PDAPP/WT brain (Fig. 4B). Cell counts showed a relative loss of astrocytes and endothelial cells, and a concomitant increase in microglia and macrophages, in aged PDAPP/KO compared to PDAPP/WT brain regions (Fig. 4C). A relative increase in microglia and macrophages was already evident in PDAPP/KO females at 20 weeks of age (Fig. 4D), indicating expansion of inflammatory cell types to precede amyloid and tau pathologies, as well as associated cell death seen at 40 weeks of age. No change in relative numbers of microglia and macrophages, or other sorted cell types, was seen in cortex or hippocampus of 40 weeks old female KO lacking PDAPP when compared to WT controls (Fig. 4E-F).

As SORCS2 deficiency increased the sensitivity of primary astrocytes to Aβ stress *in vitro*, we focused further analyses on this cell type in PDAPP/KO mice. Immunostaining for astrocyte markers astrocyte cell surface antigen-2 (ASCA2) and S100 calcium binding protein B (S100β) confirmed an overall reduction in astrocyte immunoreactivity in aged PDAPP/KO female brains (Fig. 4G). To interrogate the astrocyte subpopulation lost, these cells were analyzed by flow cytometry using a combination of markers GFAP, aldehyde dehydrogenase 1 family member L1 (ALDH1L1), S100β, and ACSA2 (Fig. S4A). Reduced cell numbers were observed for all astrocyte subpopulations with exception of the S100β^+^/ACSA2^+^ fraction, which includes astrocytes but also oligodendrocytes (Fig. S4B). While PDAPP/KO brains contained fewer GFAP^+^ cells, GFAP levels in the remaining astrocytes were increased compared to PDAPP/WT brains as shown by flow cytometry (Fig. 4H). An increase in GFAP levels in remaining astrocytes was also documented by Western blot (Fig. 4I) and immunohistology (Fig. 4J-K), arguing for astrogliosis in aged PDAPP/KO. Astrogliosis was not seen in 40 weeks old female KO mice lacking PDAPP, as deduced from brain GFAP levels being comparable to that in matched WT (Fig. S4C).

### Aβ induces aggravated microglia activation and pro-inflammatory responses in SORCS2-deficient mice

Activation of microglia and macrophages represents an important aspect of the inflammatory brain response to amyloid deposits (reviewed in (*27, 28*)). FACS documented an overall increase in inflammatory cell types in aged PDAPP/KO female brains with a relative shift from microglia to macrophages (Fig. 5A). This phenotype was already observable in 20 weeks old PDAPP/KO females (Fig. 5B), but not in 40 weeks old KO animals lacking PDAPP (Fig. 5C). These data substantiated Aβ-induced changes in inflammatory cell type composition as an early feature of SORCS2-deficient brains. Resident microglia and macrophages in PDAPP/KO brains were characterized by an increase in IBA1 expression as shown by immunohistology (Fig. 5D) and Western blot (Fig. 5E). A shift towards a pro-inflammatory profile was supported by reduced transcript levels for *Cd163* and *Cd206*, characteristic of anti-inflammatory microglia (Fig. 5F) (*29, 30*). Induction of an inflammatory brain milieu in PDAPP/KO mice was further corroborated by cytokine profiling, documenting an increase in pro-inflammatory cytokines and chemokines (Suppl. Table 1). These changes were most obvious in the cohort of young PDAPP/KO mice with low Aβ levels, and seen to a lesser extent in young or aged PDAPP/KO animals with high brain Aβ load. These findings further argued that a pro-inflammatory milieu in PDAPP/KO brains was an early feature of SORCS2 deficiency, preceding full-blown amyloid and tau co-morbidity observed at a later stage.

**Figure 5:**
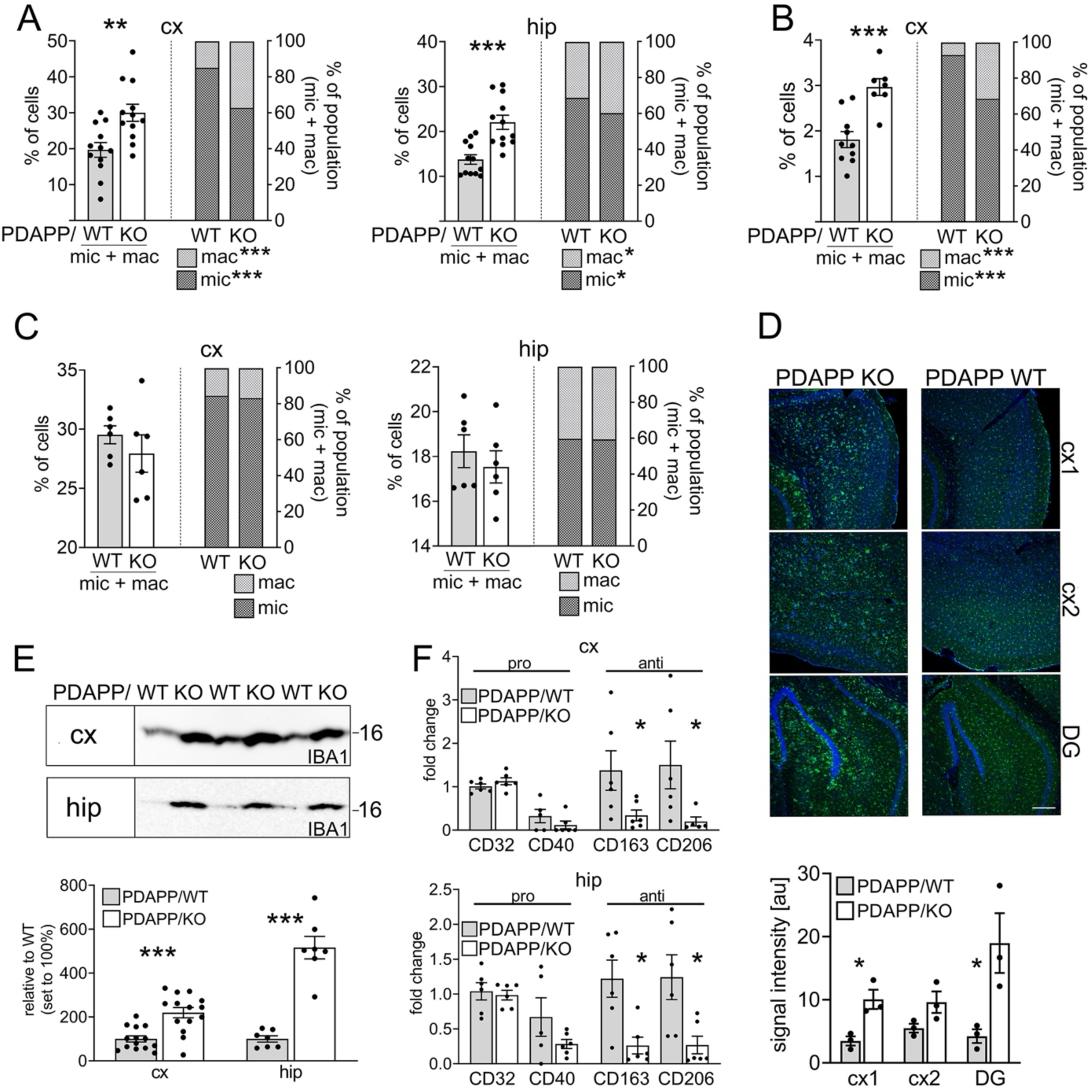
Enhanced pro-inflammatory microglia response in AD mice lacking SORCS2 **(A)** Quantification of cell numbers of microglia and macrophages in cortex (cx) and hippocampus (hip) of 40 weeks old PDAPP/WT and PDAPP/KO mice using FACS. Data are expressed as % of the total number of cells (left axis) or as % of the microglia and macrophage subpopulation (right axis), and are given as mean ± SEM from n=12 animals per genotype (two-sided unpaired Student’s *t*-test). PDAPP/KO females exhibit a significant increase in combined cell numbers with a relative shift from microglia to macrophages in both brain regions. (**B**) Experiment as in A, but quantifying the number of microglia and macrophages in cx of 20 weeks old PDAPP/WT and PDAPP/KO female mice. Data are given as mean ± SEM (n=9-11 animals per genotype, unpaired Mann-Whitney U test). **(C)** Experiments as in A, but using brain samples from 40 weeks old WT and KO mice lacking PDAPP. Data are given as mean ± SEM from n=6 animals per genotype. No significant differences in cell numbers are seen comparing genotypes (unpaired Mann-Whitney U-test). **(D, upper panel)** Representative immunofluorescence images of cortical regions (cx1, cx2) and the dentate gyrus (DG) of PDAPP/KO and PDAPP/WT females stained for IBA1 (green). Nuclei were counterstained with DAPI (blue). Scale bar: 200 μm. **(D, lower panel)** Signal intensities for IBA1 (arbitrary units, au), as determined from immunostainings in D, indicate increased levels of IBA1 immunoreactivity in the various brain regions of aged PDAPP/KO as compared to control females. Data are given as mean ± SEM from n=3 animals per genotype (two-sided unpaired Student’s *t*-test). **(E)** IBA1 in cx and hip lysates, as determined by Western blot (upper panel), and densitometric scanning of replicate blots (lower panel), show increased levels in aged PDAPP/KO compared to PDAPP/WT females. Data are expressed as relative to WT (set to 100%) and given as mean ± SEM from n=7-14 animals per genotype (two-sided unpaired Student’s *t*-test for cortex, unpaired Mann-Whitney U test for hippocampus). **(F)** Transcript levels for markers of pro-or anti-inflammatory responses in microglia sorted from cx and hip of 40 weeks old PDAPP/WT and PDAPP/KO female mice. Data are expressed as relative to WT (set to 1) and given as mean ± SEM for n=6 animals per genotype (unpaired Mann-Whitney U-test). *, P < 0.05; **, P < 0.01; ***, P < 0.001

### Astrocyte-specific loss of SORCS2 recapitulates features of global receptor deficiency

To dissect cell-type specific actions of SORCS2 in the context of Aβ stress, we generated novel mouse models with conditional inactivation of *Sorcs2* in either neurons or astrocytes. In detail, we crossed mice homozygous for the floxed *Sorcs2* allele (loxWT) (*16*) with transgenic lines constitutively expressing Cre recombinase under control of the pan-neuronal *Baf53b* promoter (31) or carrying a tamoxifen-inducible Cre-ERT2 transgene driven by the astrocyte-specific ALDH1L1 promoter (*32*). Both (loxWT x Cre) lines were bred with the PDAPP strain and referred to as neuron-specific (PDAPP/nsKO) or astrocyte-specific (PDAPP/asKO) lines. Western blot analysis documented a significant reduction in SORCS2 levels in cortical and hippocampal extracts from PDAPP/asKO as compared to PDAPP/loxWT female mice at 38 weeks of age (Fig. S5A). Using qRT-PCR on sorted cells, loss of expression was attributed to reduced *Sorcs2* transcript levels in cortical and hippocampal astrocytes, but not in neurons (Fig. S5B). Astrocyte-specific loss of *Sorcs2* transcript and protein levels could also be induced in younger mice at 30 weeks of age (Fig. S5C-D). In young PDAPP/asKO mice, soluble Aβ levels initially decreased in the hippocampus when compared to matched PDAPP/loxWT (Fig. 6A), mirroring the phenotype seen in primary astrocytes in Fig. 1B above. This trend reversed in aged PDAPP/asKO animals when hippocampal levels of Aβ_40_ and Aβ_42_increased compared to age-matched PDAPP/loxWT (Fig. 6B), recapitulating the impact of global SORCS2 deficiency in aging PDAPP/KO mice. These distinct effects of astrocytic SORCS2 deficiency on Aβ levels in hippocampi were less pronounced in the cortices of PDAPP/asKO females (Fig. 6A-B), possibly reflecting differences in efficiency of *Sorcs2* inactivation or astrocyte vulnerability in the two brain regions.

**Figure 6:**
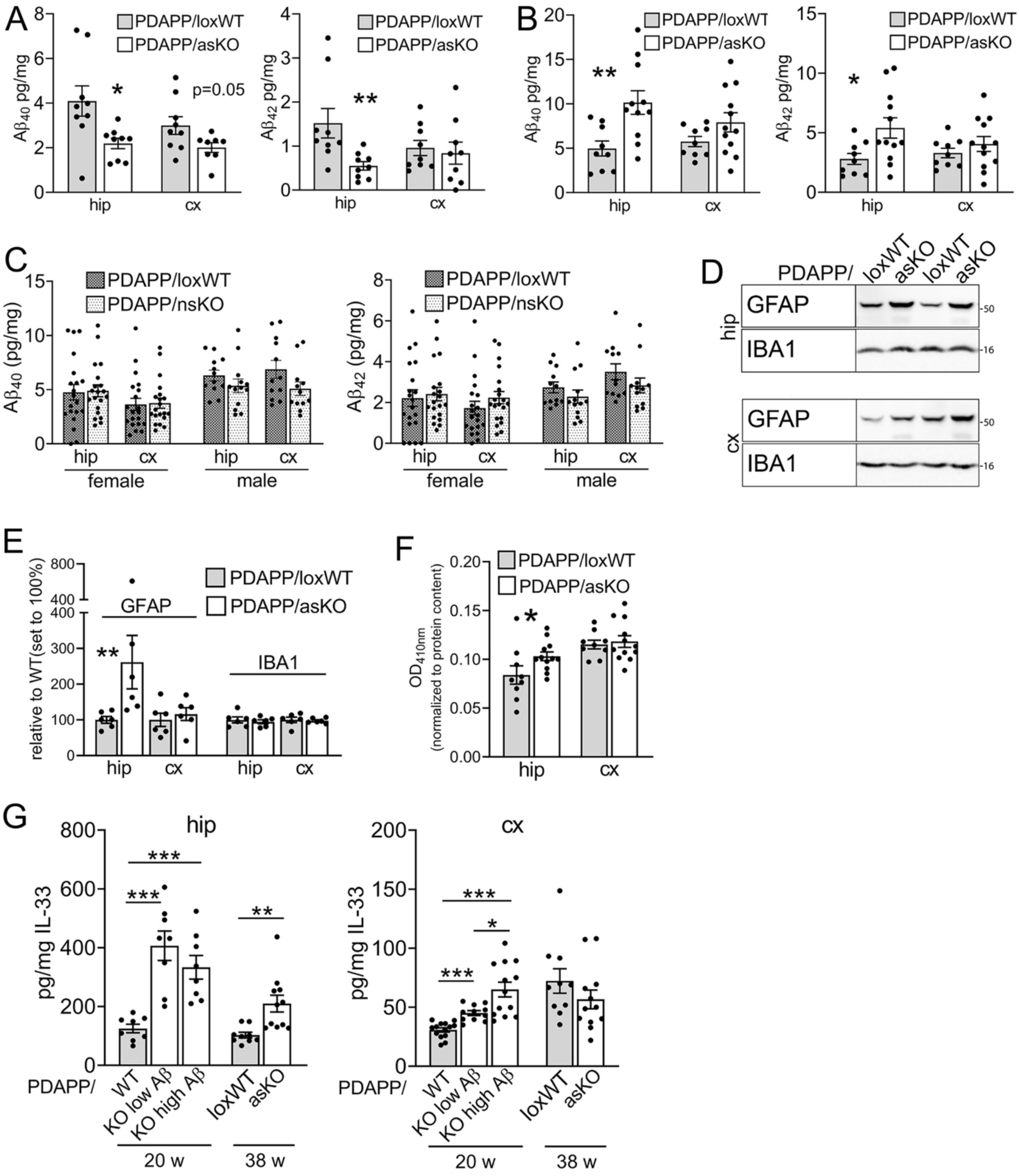
Astrocyte-specific inactivation of *Sorcs2* increases Aβ levels and induces cell stress in the AD mouse brain (**A**) Levels of soluble Aβ_40_ and Aβ_42_ in hippocampus (hip) and cortex (cx) of 32 weeks old PDAPP animals (20 weeks post tamoxifen injection) homozygous for *Sorcs2^lox/lox^* (PDAPP/loxWT) or carrying an astrocyte-specific *Sorcs2* gene defect (PDAPP/asKO). Data are given as mean ± SEM from n=9 animals per genotype (two-sided unpaired Student’s *t*-test). **(B)** Data as in A, but from hip and cx extracts of 38 weeks old PDAPP/loxWT and PDAPP/asKO animals. Data are given as mean ± SEM for n=9-12 animals per genotype (two-sided unpaired Student’s *t*-test). **(C)** Levels of soluble Aβ_40_ and Aβ_42_ in cx and hip of 38 weeks old male or female PDAPP mice, either PDAPP/loxWT or carrying a neuron-specific *Sorcs2* gene defect (PDAPP/nsKO). Data are given as mean ± SEM for n=16-20 animals per genotype (unpaired Mann-Whitney U-test). **(D-E)** Representative Western blot for GFAP and IBA1 in hip (D, upper panel) or cx brain extracts (D, lower panel), and quantification from densitometric scanning of replicate blots thereof (E), are shown. Hip GFAP levels increase in 38 weeks PDAPP/asKO females compared to matched PDAPP/loxWT controls. Data are expressed as relative to WT (set to 100%) and given as mean ± SEM from n=6 animals per genotype (unpaired Mann-Whitney U-test). **(F)** Nucleosome fragmentation assay documents increased cell death in hip extracts of 38 weeks old PDAPP/asKO females compared to matched PDAPP/WT. Data are given as mean ± SEM from n=9-12 animals per genotype (unpaired Mann-Whitney U-test). **(G)** Levels of IL33 in hip and cx extracts of PDAPP/WT or PDAPP/KO female mice at 38 weeks of age. Young females are separated into two groups based on low versus high brain Aβ levels (cut off: 300 pg/ml Aβ_40_ and 800 pg/ml Aβ_42_ for hip; 35 pg/ml Aβ_40_ and 60 pg/ml Aβ_42_ for cx). Data are given as mean ± SEM from n=8-14 animals per genotype (two-sided unpaired Student’s *t*-test) *, P < 0.05; **, P < 0.01; ***, P < 0.001

Loss of SORCS2 expression was also seen in cortices of PDAPP/nsKO females (Fig. S5E). However, *Sorcs2* transcript levels were reduced in both astrocytes and neurons in these animals (Fig. S5F). By contrast, neuron-specific *Sorcs2* inactivation was achieved in males as documented by a significant reduction in cortical protein levels (Fig. S5E) that resulted from a decrease in *Sorcs2* transcripts in sorted neurons but not astrocytes (Fig. S5F). Still, neuron-specific loss of SORCS2 did not impact cortical or hippocampal levels of Aβ in aged PDAPP/nsKO males compared to matched PDAPP/loxWT (Fig. 6C). Recapitulating additional features of global SORCS2 deficiency, astrocyte-specific loss of SORCS2 increased GFAP levels (Fig. 6D-E) and induced cell death (Fig. 6F) in hippocampi of aged PDAPP/asKO females. Also, levels of the early stress marker IL33 were significantly increased in cortex and hippocampus. This effect was more pronounced in young PDAPP/KO mice with modest Aβ load than in young or aged PDAPP/asKO with higher Aβ levels, when compared to their respective age-matched genotype controls (Fig. 6G).

### Aβ stress in astrocytes results in aberrant induction of δ secretase activity

Taken together, our studies in obligate and conditional SORCS2-deficient mouse models argued for loss of receptor activity in astrocytes to induce AD-like pathologies. Because these phenotypes required an initial trigger from the PDAPP transgene, we focused on amyloidogenic processes as possible cause of these defects. As most dramatic phenotypes were seen with global SORCS2 deficiency, we carried out these studies in PDAPP/KO mice.

Initially, we measured levels of soluble (s) APPα and sAPPβ, cleavage products of APP processing by α and β secretases, respectively (Fig. 7A). Surprisingly, levels of sAPPα were reduced, but levels of sAPPβ were undetectable in cortex or hippocampus of 40 weeks old PDAPP/KO females when compared to PDAPP/WT animals (Fig. 7B). An explanation for this apparent paradox of massive production of Aβ, albeit at undetectable levels of sAPPβ, may stem from the activity of asparagine endopeptidase (AEP), also known as legumain or δ secretase.

**Figure 7:**
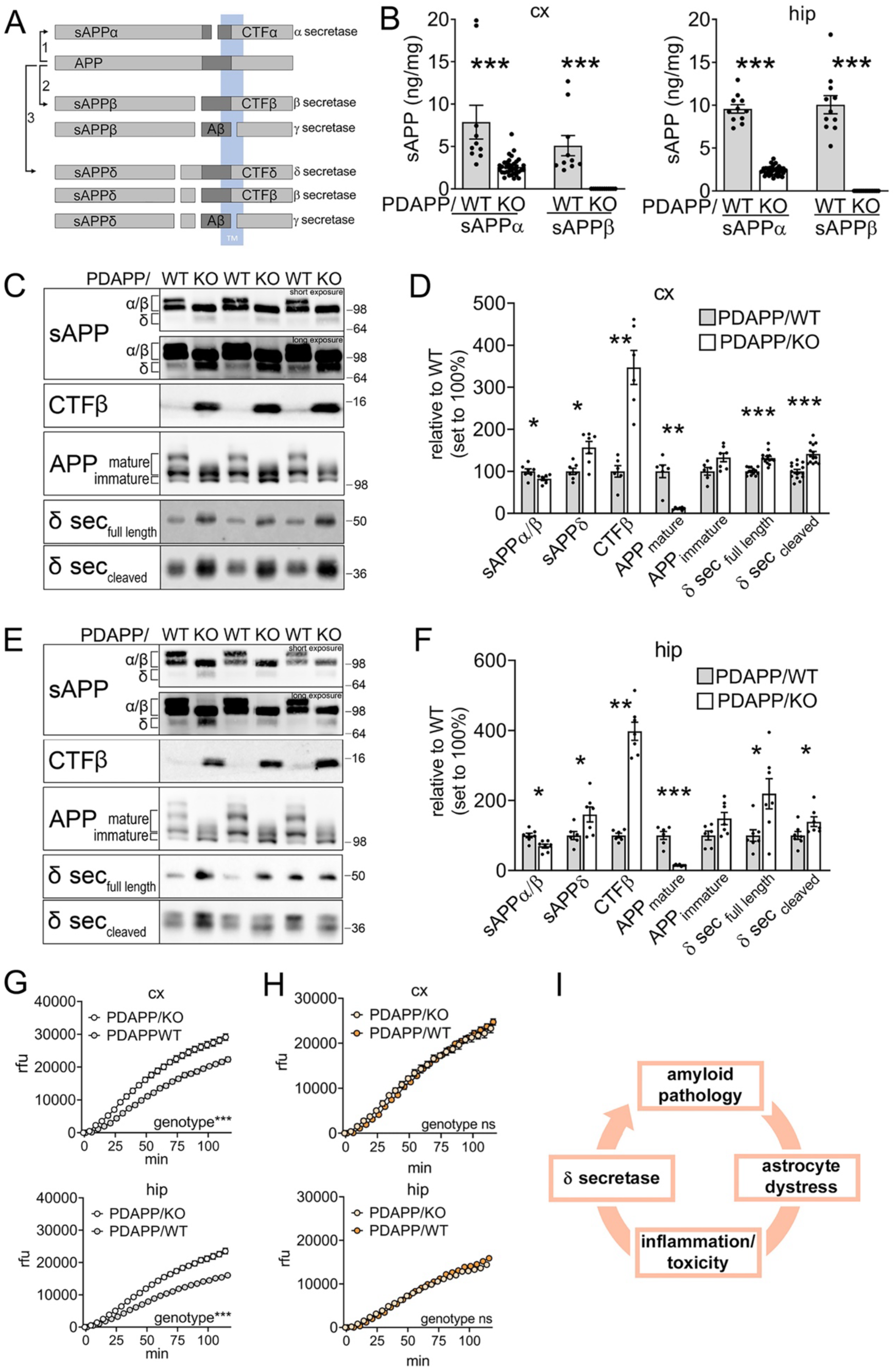
Amyloid peptide overproduction in SORCS2-deficient mice is caused by enhanced δ secretase activity (**A**) Schematic presentation of APP processing pathways and the resulting cleavage products. (1) non-amyloidogenic processing initiated by α secretase, (2) amyloidogenic processing by β and γ secretases, (3) alternative amyloidogenic processing by sequential cleavage from δ, β, and γ secretases. CTFδ and sAPPδ are cleavage products unique to δ secretase action. (**B**) Levels of soluble (s)APPα and sAPPβ in cortex (cx) and hippocampus (hip) of 40 weeks old female PDAPP/WT and PDAPP/KO animals as determined by ELISA. Levels of sAPPα are reduced, while levels of sAPPβ are undetectable in PDAPP/KO. Data are given as mean ± SEM from n=11-36 animals per genotype (unpaired Mann-Whitney U test for cx, two-sided unpaired Student’s *t*-test for hip). **(C, D)** Representative Western blot (C), and densitometric scanning of replicate blots (D), quantifying levels of APP, CTFβ, sAPP species, as well as full-length and cleaved forms of δ secretase in the cx lysates of 40 weeks old PDAPP/WT and PDAPP/KO females. Immunoreactive bands representing mature and immature forms of APP as well as APPα/b and sAPPδ are marked. Data are expressed as relative to WT (set to 100%) and given as mean ± SEM from n=6-7 (for APP, CTFβ, and sAPPs) and n=14 (for δ secretase) animals per genotype (unpaired Mann-Whitney U test for APP, CTFβ, and sAPP; two-sided unpaired Student’s *t*-test for δ secretase). **(E, F)** Experiment as in C and D, but testing hip extracts of 40 weeks old PDAPP females. **(G)** Activity of δ secretase is increased in cx and hip lysates of 40 weeks old PDAPP/KO compared to PDAPP/WT females, as documented by activity assays using a fluorogenic substrate (see methods for details). Data are given as mean ± SEM from n=6-7 animals per genotype (repeated measures two-way ANOVA to test significance for genotypes). **(H)** Experiment as in G, but using cx and hip extracts from 12 weeks old PDAPP/WT and PDAPP/KO females, showing comparable levels of δ secretase activity. Data are given as mean ± SEM from n=3 animals per genotype (repeated measures two-way ANOVA). rfu, relative fluorescence units. *, P < 0.05; **, P < 0.01; ***, P < 0.001. (**I**) Model of pathological cascade induced by Aβ stress imposed on astrocytes (see discussion section for details).

Expression of δ secretase is induced during acute or chronic insults to the brain, as in AD, Parkinson’s disease, stroke, or epilepsy (*33–36*). The enzyme acts on APP, generating an extended stub (called CTFδ in Fig. 7A) that represents a favorable substrate for β secretase processing. Cleavage by δ secretase generates a shortened sAPP fragment (sAPPδ) that presumably lacks the recognition site for antibodies used in commercial sAPPβ ELISA (www.mesoscale.com), providing an explanation for our inability to detect sAPPβ in PDAPP/KO mice. This assumption was substantiated by Western blotting showing a shorter sAPPδ fragment in cortical (Fig. 7C-D) and hippocampal (Fig. 7E-F) extracts from aged PDAPP/KO compared to PDAPP/WT females. Aggravated amyloidogenic processing was substantiated by an increase in CTFβ stubs, and a concomitant depletion of mature APP from cortex (Fig. 7C-D) and hippocampus (Fig. 7E-F) of PDAPP/KO mice.

Induced expression of δ secretase as likely cause of aberrant amyloidogenic processing in PDAPP/KO was supported by elevated levels of full-length (pro-form) and cleaved (active) forms of the enzyme in cortex (Fig. 7C-D) and hippocampus (Fig. 7E-F). Ultimately, overactivity of δ secretase in cortex and hippocampus of aged PDAPP/KO mice was confirmed using an activity assay based on proteolytic cleavage of a fluorogenic substrate (Fig. 7G, see methods). Increased level (Fig. S6A) and activity (Fig. S6B) of δ secretase was also seen in cortex and hippocampus of aged PDAPP/KO males. Contrary to aged PDAPP/KO animals,

δ secretase activity was not increased in young PDAPP/KO mice at 12 weeks of age, indicating that induction of δ secretase activity in SORCS2-deficient brains aligns with appearance of profound amyloid pathology (Fig. 7H). Neither levels of murine APP (Fig. S6C) nor levels (Fig. S6D) or activity (Fig. S6E) of δ secretase were altered in aged KO female mice lacking PDAPP, corroborating that SORCS2 deficiency alone is insufficient to induce non-canonical APP processing.

## DISCUSSION

Mechanisms defining the role of astrocytes in age-related dementia still remain poorly understood. Now, our data identified astrocyte distress as a central disease-promoting process at early stages of AD (Fig. 7I). According to our concept, the inability to cope with aggravated insults from Aβ results in astrocyte reactivity, triggering widespread gliosis and brain inflammation. These stress responses, in turn, induce δ secretase, accelerating amyloidogenic processing and tau pathology, and resulting in a vicious cycle that drives exhaustion and eventual death of astrocytes.

Amyloid plaques and tau hyperphosphorylation are two distinguishing features of AD in patients. An unprecedented comorbidity, characterized by massive increase in αβ and murine tau hyperphosphorylation, is seen in a mouse model of hypersensitivity of astrocytes to Aβ, induced by inactivation of the protective stress response gene *Sorcs2*. Increased astrocyte distress coincides with a pro-inflammatory brain milieu and with prominent gliosis, further characteristics of human AD. These phenotypes depend on a trigger from PDAPP. The extent of amyloid and tau comorbidities in PDAPP/KO mice is striking, given that *APP^Ind^* is a rather subtle murine model of AD characterized by modest levels of human Aβ and limited plaque deposition visible only after 9 months of age (*15*). Thus, aggravation of phenotypes with a 30-fold increase in Aβ_42_ levels and tau hyperphosphorylation, in the absence of a human *TAU* transgene, clearly stems from the lack of protective SORCS2 actions.

The temporal succession of phenotypic features seen with various mouse genotypes and ages clearly establishes the causality of pathological processes related to astrocyte dysfunction in our AD model. Pro-inflammatory activation of glia, as evidenced by cytokine profiling, is a phenotype already obvious in young PDAPP/KO mice with low Aβ burden (Table S1). A relative increase in immune cell types (Fig. 4D), with a shift from microglia to macrophages (Fig. 5B) corroborates a pro-inflammatory milieu in young PDAPP/KO brains as an early consequence of SORCS2 deficiency, preceding full-blown amyloid and tau pathologies at later stages. This conclusion is supported by induction of IL33, an early indicator of cell stress (Fig. 6G). Increased Aβ levels are also seen in aged mice with astrocytic *Sorcs2* inactivation (Fig. 6B). Although less pronounced than in the obligate PDAPP/KO model, likely due to technical limitations of conditional receptor depletion, these findings argue that astrocytic SORCS2 deficiency plays a decisive role in AD pathology.

Prior studies have confirmed the sensitivity of astrocytes to Aβ exposure. Astrocytes participate in removal of amyloid from the brain parenchyma (*37, 38*). However, contrary to other cell types, such as microglia, astrocytes accumulate rather than degrade ingested material (*39, 40*). As a consequence, they acquire a high intracellular load of toxic amyloid that impacts endo-lysosomal function and energy homeostasis (*3, 39, 41*). Exposure to Aβ also triggers release of pro-inflammatory cytokines and chemokines from astrocytes (*10, 42, 43*), evoking inflammation that further enhances brain Aβ production (*44–46*) and tau pathology (*45*). SORCS2-deficient astrocytes exhibit increased uptake and intracellular accumulation of Aβ (Fig. 1B-D). Amyloid buildup coincides with astrocyte reactivity (Fig. 1E), defects in lysosomal acidification (Fig. 1F), and apoptotic cell death (Fig. 1G-I). These findings document a role for SORCS2 in preventing excessive uptake and/or accumulation of Aβ in astrocytes. Enhanced clearance of Aβ by SORCS2-deficient astrocytes is supported by reduced Aβ levels initially seen in the brains of PDAPP /asKO animals before progressing pathology increases amyloid burden (Fig. 6A).

SORCS2 is an intracellular sorting receptor that acts in protective stress responses in multiple tissues. In pancreatic alpha cells, it directs secretion of osteopontin, a factor that stabilizes insulin release from beta cells under glucose stress (*47*). In neurons, SORCS2 sorts the amino acid transporter EAAT3 to the cell surface, increasing cysteine import and glutathione production as means of oxidative stress response in epilepsy (*48*). In astrocytes, the receptor facilitates release of endostatin to promote angiogenesis in the post-stroke brain (*13*). Given the multifunctionality of SORCS2 (*12*), proteins sorted by this receptor in astrocytes are difficult to predict but may include clearance receptors for Aβ. One possible candidate is the neurotrophin receptor p75NTR, an established interacting partner of SORCS2 (*14, 49*) implicated in Aβ binding and uptake (*50, 51*). Potentially, SORCS2-dependent sorting of p75NTR reduces Aβ ingestion, protecting astrocytes from amyloid overload. In line with this hypothesis, the closely related VPS10P domain receptor SORCS3 was shown to target p75NTR to lysosomes to promote its degradation (*52*). Whatever the protective function of SORCS2 in the context of AD may be, it is universal to the entire astrocyte lineage as concluded from loss of all tested astrocyte subtypes in the PDAPP/KO brain (Fig. S4B).

Astrocyte reactivity has been proposed to enhance amyloid burden, yet the underlying molecular mechanism remained elusive (reviewed in (*53*)). Our findings shed light on this process by identifying a surprising culprit, δ secretase. This endolysosomal peptidase cleaves APP at N585 and N373, accelerating Aβ production by generating a favored β secretase substrate (*34, 54*).

Remarkably, δ secretase activity also drives tau pathology as it proteolytically inactivates I2PP2A, an inhibitor of PPA2, the main enzyme responsible for dephosphorylation of tau (*55–57*). Consequently, δ secretase actions result in tau hyperphosphorylation and aggregation, accelerating AD pathology in mouse models double transgenic for human *APP* and *TAU* (*54, 58*). Expression of δ secretase increases with brain age (*34*) and is seen most highly in microglia and astrocytes under acute or chronic distress (*36*). With relevance to our hypothesis, δ secretase expression and activity in the brain is induced by inflammation (*59, 60*) and by lysosomal substrate overload (*61*), two cellular phenotypes prevalent in SORCS2-deficient astrocytes.

Whatever the exact trigger(s) in our mouse model may be, our findings identify induction of δ secretase activity as a molecular mechanism whereby astrocyte stress provides an early trigger for amyloid and tau comorbidities in the AD brain.

## ACKNOWLEDGEMENT

The authors are grateful to K. Kampf, T. Pasternack, and R. Vogel for expert technical assistance.

## Funding

National Science Center OPUS program, 2020/37/B/NZ3/00761 (ARM)

Research University Program at the University of Warsaw I.3.4 Action of the Excellence Initiative (ARM)

Alzheimer Forschung Initiative #23001R (TEW) Novo Nordisk Foundation NNF18OC0033928 (TEW)

## Author contributions

VS, ARM, and TEW conceptualized the study. VS, EZ, TO, EZ-P, JC, and ARM performed experiments and evaluated data. BLH contributed essential mouse lines. TEW wrote the manuscript with help from VS and ARM.

## Data and materials availability

All data and materials described in this study are available upon reasonable request from the corresponding authors of this study.

## SUPPLEMENTARY MATERIALS

Supplementary Methods

Supplementary figure legends

Figs. S1 to S5

Table S1

