## Supplemental methods, figures, and tables for "Aβ-induced distress of astrocytes triggers Alzheimer disease pathology through non-canonical δ secretase activity"

### SUPPLEMENTARY METHODS

#### Expression analyses in mouse tissues and primary cell types

Protein extracts from cortical or hippocampal brain tissues were generated using standard protocols. All steps were performed at 4°C using solutions containing protease (Complete #11836145001, Roche) and phosphatase (PhosStop 04906837001, Roche) inhibitors. Soluble and membrane protein fractions were obtained by homogenization of brain samples in solution A (20 mM Tris-HCl, 2 mM $MgCl_2$, 250 mM sucrose, pH 7.5) using a pestle B (10 strokes). After centrifugation at 1000xg for 5 min, the supernatants were transferred to an ultracentrifuge tube. The remaining pellet was again homogenized in solution A and centrifuged, and both supernatants combined. Soluble protein fractions were obtained by ultracentrifugation of the combined supernatants at 175,000xg for 30min. The remaining pellets were lysed in lysis buffer (20 mM Tris-HCl, 10 mM EDTA, 1% NP40, 1% Triton X-100, pH 7.4) to derive the membrane protein fraction. Total protein extracts were obtained by homogenization of brain tissues in lysis buffer directly.

For Western blotting, protein extracts from mouse brain tissues or primary astrocyte cultures were subjected to SDS-PAGE in Tris-glycine gels, followed by transfer to nitrocellulose membranes. For immunodetection, the following primary antibodies (diluted in 20 mM Tris, 150 mM NaCl, 0.1% Tween-20, 5% BSA) were used: APP antiserum (1227, in-house), δ-secretase (Legumain #93627, Cell Signaling Technology), SORCS2 (AF4237, R&D Systems), IBA1 (ab5076, Abcam), GFAP (G3893, Sigma Aldrich; #20334, DAKO Agilent), cleaved caspase-3 (#9664, Cell Signaling Technology), cleaved PARP (Asp214, #9541, Cell Signaling Technology) tau (HT7, Invitrogen #MN1000), ptau (AT8, Invitrogen #MN1020; AT180; Invitrogen # MN1040) or beta actin (Abcam #ab8227).

For ELISA, soluble protein fractions of brain extracts were used to measure various analytes. APP processing products were determined by multiplex assays (V-PLEX Plus Aβ Peptide Panel 1 (4G8), #K15199G; sAPPα/sAPPβ Kit, #K15120E; Meso Scale Discovery). Levels of IL-1β, TNFα, IL-12p70, IL-6, IP10/CXCL10, IL-33, MIP1α/CCL3, IL-2, IL-5, IL-16, IL-10, IL-4, and IL-22 were determined by custom-made multiplex assays using the Meso Scale Discovery protocol. TGF-β2 /TGF-β3 levels were determined using the U-PLEX TGF-β Combo multiplex assay (#K15242K, Meso Scale Discovery). Phospho and total tau levels were measured using Kit #K15121D (Meso Scale Discovery). PikoKine ELISA was used to quantify YKL40/Chi3l1 and IL33 (Boster Biological Technology #EK0975, #EK0930) according to the manufacturer's protocols.

#### Quantitative RT-PCR

Total RNA was extracted for brain tissues using the RNeasy Plus Micro Kit (Qiagen # 74034) combined with the RNase-free DNase Set (Qiagen # 79254) or from primary astrocyte cells using the Direct-zol RNA Miniprep Kit (Zymo Research #R2052) according to the manufacturers' instructions. The RNA was converted to cDNA (Applied Biosystems # 4387406; #4368814) and gene expression analysis (Applied Biosystems # 4369016; #4366072) was performed using the following probes: *Sorcs2* (Mm00473050_m1 and Mm01217942_m1), *Iba1/Aif1*(Mm00479862_g1), *Tmem119* (Mm00525305_m1), *Cdh5* (Mm00486938_m1), *Vwf* (Mm00550379_m1), *Mbp* (Mm01266402_m1), *Mog* (Mm01273867_m1), *Gfap* (Mm01253033_m1), *Aldh1l1* (Mm03048949_m1), *NeuN/Rbfox3* (Mm01248771_m1), *Baf53b/Actl6b* (Mm00504274_m1), *Cd32* (Mm 00438875_m1), *Cd40* (Mm 00441891_m1), *Cd163* (Mm 00474091_m1), *Cd206* (Mm 01329359_m1), *Rn18s* (Mm03928990_g1) (Thermo Fisher Scientific).

**Immunohistochemistry**

Mice were perfused and fixed with 4% paraformaldehyde in PBS. Brains were carefully dissected and postfixed for additional 24 h before treatment in 30% sucrose/PBS for several days at 4°C. Free-floating 50 μm sections were processed by 10 min antigen retrieval in 10 mM citric acid/TBA (20mM Tris, 150mM NaCl pH 6.0) at 80°, followed by 10 min permeabilization in PBS/0.05% Tween-20 with 0.3% Triton X-100, and blocking with M.O.M. (Vector Labs #MKB-2213) for 60 min. In case of GFAP and IBA1 stainings, antigen retrieval and permeabilization were not performed. Next, 40 μm free-floating sections were blocked for 1h in 1% horse serum in PBS. For immunodetection, the sections were incubated overnight at 4°C with primary antibodies diluted in incubation buffer (PBS with 1% bovine serum albumin, 1% normal donkey serum, 0.3% Triton X-100). Then, sections were washed in PBS and treated for 2 hours at room temperature with fluorochrome-conjugated secondary antibodies (Alexa, Invitrogen). After washing in DAPI, stained sections were mounted in DAKO Fluorescence Mounting Medium (F4680, Sigma Aldrich). For immunodetection, the following primary antibodies were used: tau (HT7, Invitrogen #MN1000; cleaved tau (tauC3), Invitrogen #AHB0061), ACSA2 (Miltenyi Biotec #130-116-245), S100β (Novus Biologicals #NBP2-45267PE), GFAP-Cy3 (Sigma #C9205), as well as IBA1 (Wako #019-198741). Quantification of the immunosignals was done in ImageJ using the SUM option for the z-stacks at 2 μm distances. Biotinylated phospho-tau antibodies AT8, Ser202, and Thr205 (Invitrogen #MN1020B) were used to visualize tau. Quantification was performed using the Colour Deconvolution with H DAB function in ImageJ. For visualization of amyloid deposits, mouse brains were fixed in 4% paraformaldehyde/PBS for 24 hours. After dehydration, tissues were embedded in paraffin and sectioned on Super Frost Plus glass slides at 5 μm. Sections were deparaffinized and rehydrated with a series of Roti-Histol and ethanol, and stained with 1% aqueous Thioflavin-S solution (Sigma #T1892) for 8 min at room temperature. Thereafter, the sections were washed twice in 80% and once in 95% ethanol, followed by three washes in distilled water.

**Flow cytometric analysis and quantification of brain cell types**

Brain regions were dissected, gently minced, and resuspended in Hanks´ Balanced Salt Solution (HBSS, Gibco #14185-045) containing 8 units of papain (Worthington Biochemical #LK003172) and 250 units of DNase I (Invitrogen #18047-019). Resuspended tissues were incubated in a water bath at 37°C for 20 min and gently titrated 10-fold with a 20G syringe halfway through the incubation time. After incubation, 5 ml of wash buffer (HBSS with 2 mM EDTA and 0.5% BSA) were added to the homogenates and filtered through a 70 µm mesh before cells were pelleted at 350xg for 5 min. Cell pellets were resuspended in 900 µl PBS with 0.025% BSA and 100 µl Myelin Removal Beads II (Miltenyi Biotec #130-096-433) and incubated for 15 min at 4°C. Then, cells were washed and resuspended in 2 ml of wash buffer. One ml of each suspension was placed on a LS column on the magnetic field separator. The flow through (all cells without myelinated oligodendrocytes) was collected. The LS column was washed with 4 ml wash buffer, collected in the same tube. These steps were repeated with another 1 ml aliquot of the same homogenate and both cell fractions were combined. To isolate oligodendrocytes, the LS columns were removed from the magnetic field separator and additionally loaded with 3 ml of wash buffer. The flow through was collected and the myelinated oligodendrocytes were pelleted at 350xg for 5min. The resulting pellets were resuspended in 5 ml of PBS with 20% isotonic Percoll PLUS (Millipore #E0414-250 ml) and centrifuged in swingout buckets at 310xg for 20 min to remove the myelin sheath from the oligodendrocytes. The resulting pellets were washed to remove traces of Percoll PLUS, centrifuged, and combined with the main single cell suspensions. Cells were centrifuged again, resuspended in 100 µl wash buffer containing 1:100 Mouse BD Fc Block (BD Biosiences #553141), and transferred to reaction tubes. After 15-20 min of incubation at 4°C, 100 µl of wash buffer were added containing antibodies directed against markers of microglia and macrophages (CD45-BV421; 1:200, BD Biosciences #563890), endothelia (CD49a-FITC; 1:200, Miltenyi Biotec #130-107-636), oligodendrocytes (O4-PE; 1:200, Miltenyi Biotec #130-117-357) and astrocytes (ACSA2-APC, 1:100, Miltenyi Biotec #130-116-245; S100b-PE, 1:100, Novus Bio #NBP2-45267; GFAP-BV421, 1:100, Biolegend #644710; Aldh1l1-FITC, 1:100, Novus Bio #NBP2-50045F), followed by incubation overnight at 4°C. Prior to staining for GFAP-BV421 and Aldh1l1-FITC, cells were fixed and permeabilized with Phosflow lyse/fix (BD Biosciences # 558049) and Perm Buffer III (BD Biosciences #558050) according to the manufacturers' instructions. The next day, stained cells were washed and FACS sorted using BD Aria Fusion. The FlowJo 10 Software was used for quantification of cell type numbers.

**Primary astrocyte cultures**

To derive adult primary astrocytes for Aβ uptake assays, cortices of 40 weeks old PDAPP mice were dissected and prepared as described for FACS sorting above. Cell pellets were kept on ice during preparation of the isotonic gradients by diluting Percoll (GEhealthcare #17-0891-01) in 10x PBS to a stock solution of 1.12 g/ml in 1x PBS (100% SIP). The stock solution was further diluted in PBS to obtain 70% SIP, 50% SIP, and 35% SIP. The cell pellets were resuspended in 4 ml of 35% SIP and loaded onto a gradient of 4 ml of 50% and 2 ml of 70% SIP in a 15ml Falcon tube at room temperature. The prepared tubes were centrifuged in a swingout bucket at 2000 x g with acceleration 3 and break 1 for 20 min. After centrifugation, the top myelin layer was removed and cells from the interface between 35 - 50% SIP were collected in a new 15ml Falcon tube. The isolated astrocytes were washed and resuspended in DMEM/F12 medium (Gibco #11320033) containing 10% fetal bovine serum (Life Technologies, #10270106) and 1% penicillin/streptomycin (Life Technologies, #15140-122), and seeded in 24-well culture dishes pre-coated with 0.1 mg/ml poly-D-lysine hydrobromide (Sigma #P1274). For Aβ uptake, astrocytes were washed with PBS and incubated for 2 hours with 200 µl growth medium containing 1 µM Aβ40 HiLyte-488 (Ana Spec #AS-60491-01). Next, the cells were washed three times and imaged for fluorescence signals from HiLyte-488. Thereafter, cells were trypsinized (Gibco, 0.25% trypsin/EDTA, #25200056) and washed in PBS. The suspensions were centrifuged at 5000 rpm for 5min, and the cell pellets lysed in lysis buffer (20 mM Tris-HCl, 10 mM EDTA, 1% NP40, 1% Triton X-100, pH 7.4) for 1 hour on ice. The intracellular Aβ content was quantified by ELISA (Meso Scale Discovery).

To generate primary astrocytes from neonates (P0-P2), brain tissues were dissected from mouse pups with olfactory bulbs, cerebellum, and meninges removed. Tissues were washed three times at room temperature with HBSS (Gibco #14175-095) leaving approximately 1.5 ml of buffer after the last wash. Two hundred µl of solution containing 100 mg/ml trypsin (Sigma-Aldrich #T8003) and 5 mg/ml DNase I (Sigma-Aldrich #DN25) was then added and incubated for 2 min at room temperature with brief shaking. The process was stopped by adding 5 ml of Dulbecco′s modified Eagle′s medium (4500 mg/l glucose, 1 mM sodium pyruvate) supplemented with 4 mM glutamine (Gibco #31966-021), 10% (vol/vol) FBS (Gibco #10500064), and 1% of penicillin/streptomycin (Sigma #P4333). After removing the medium until approximately 1.5 ml remained, 200µl of 5 mg/ml DNase I solution was added and the tissue was disrupted by pipetting with a Pasteur pipette. Subsequently, 10 ml of fresh medium were added and the cell suspensions centrifuged at 120xg for 10 min at room temperature. Thereafter, the cell pellet was resuspended in fresh medium (1 ml per 2 brains) and plated in a T75 flask (Sarstedt #83.3911.302) pre-coated with poly-L-lysine hydrobromide (0.1mg/ml final concentration, Sigma #P1274). At DIV2, flasks were washed three times with 1x PBS (Gibco #14200) and 15 ml of fresh cell medium were added. At DIV10, flasks were shaken on a horizontal shaker at 80 rpm and 37°C for 60 min to remove microglia population. Fresh medium was added to the remaining astrocytes. At DIV15, flasks were shaken overnight at 180 rpm, washed three times with 1X PBS, trypsinized, and cells plated on Petri dishes. At DIV20, cells were harvested and stored in liquid nitrogen. For each experiment, WT and KO astrocytes were thawed, plated, and reseeded after 3-5 days.

### SUPPLEMENTARY FIGURES AND LEGENDS

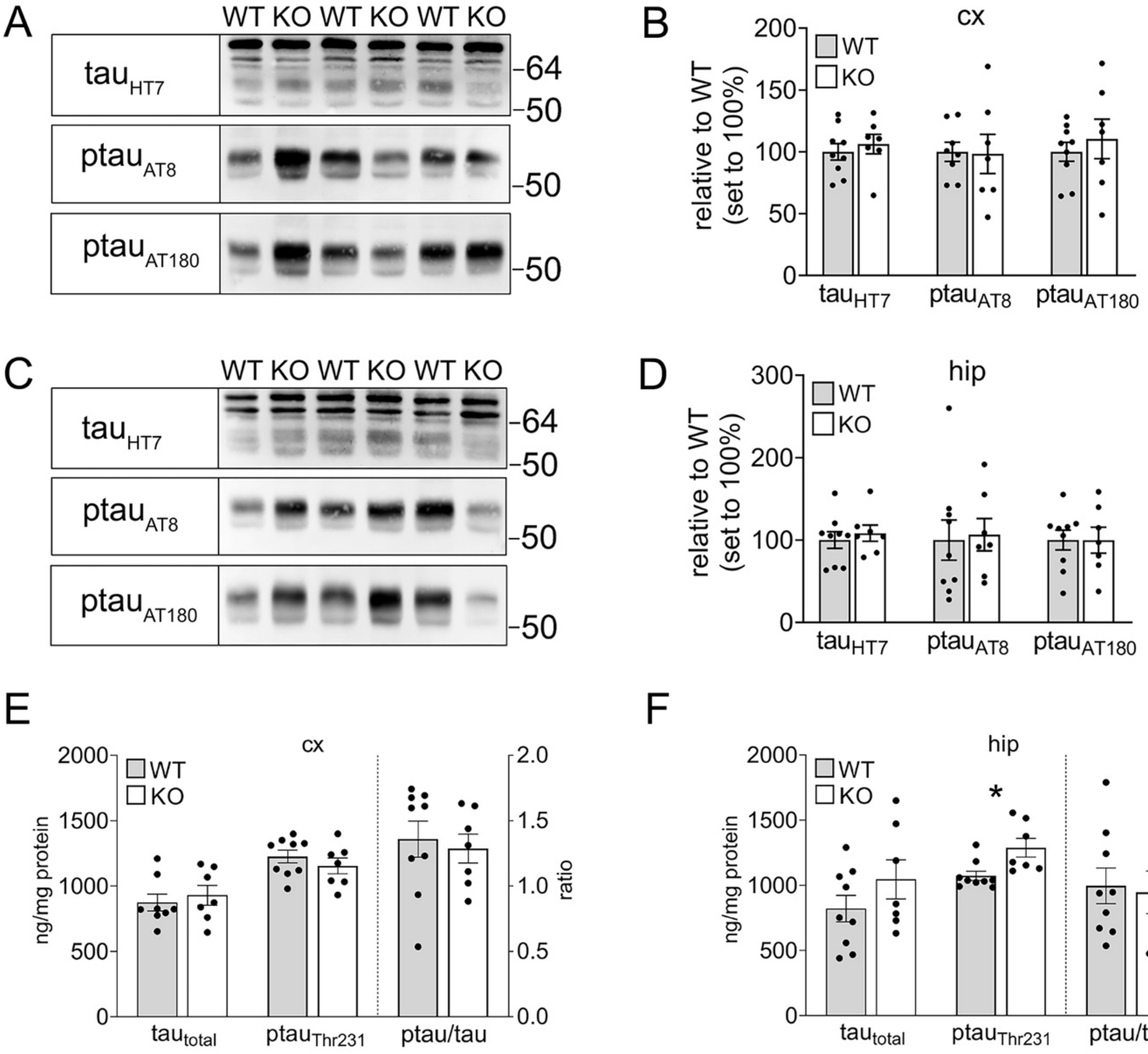


**Figure S1: Tau pathology in SORCS2-deficient female mice requires Aβ trigger**
**(A, B)** Representative Western blot analyses, and densitometric quantification of replicate blots thereof, document levels of total tau (HT7) as well as phosphorylated variants $ptau_{Ser202/Thr205}$ (AT8) and $ptau_{Thr231}$ (AT180) in cortical extracts of 40 weeks old WT and KO female animals lacking the PDAPP transgene. Data are expressed as relative to WT (set to 100%) and given as mean ± SEM from n=7-9 animals per genotype (unpaired Mann-Whitney U-test). (**C, D**) Data as in A and B but for hippocampal extracts. **(E, F)** Levels of total ($tau_{total}$) and phosphorylated variants ($ptau_{Thr231}$) of tau, as well as ratio of $ptau_{Thr231}/tau_{total}$, in cx (E) and hip (F) of 40 weeks old WT and KO females. Data are given as mean ± SEM from n=7-9 animals per genotype (unpaired Mann-Whitney U-test). *, $P < 0.05$

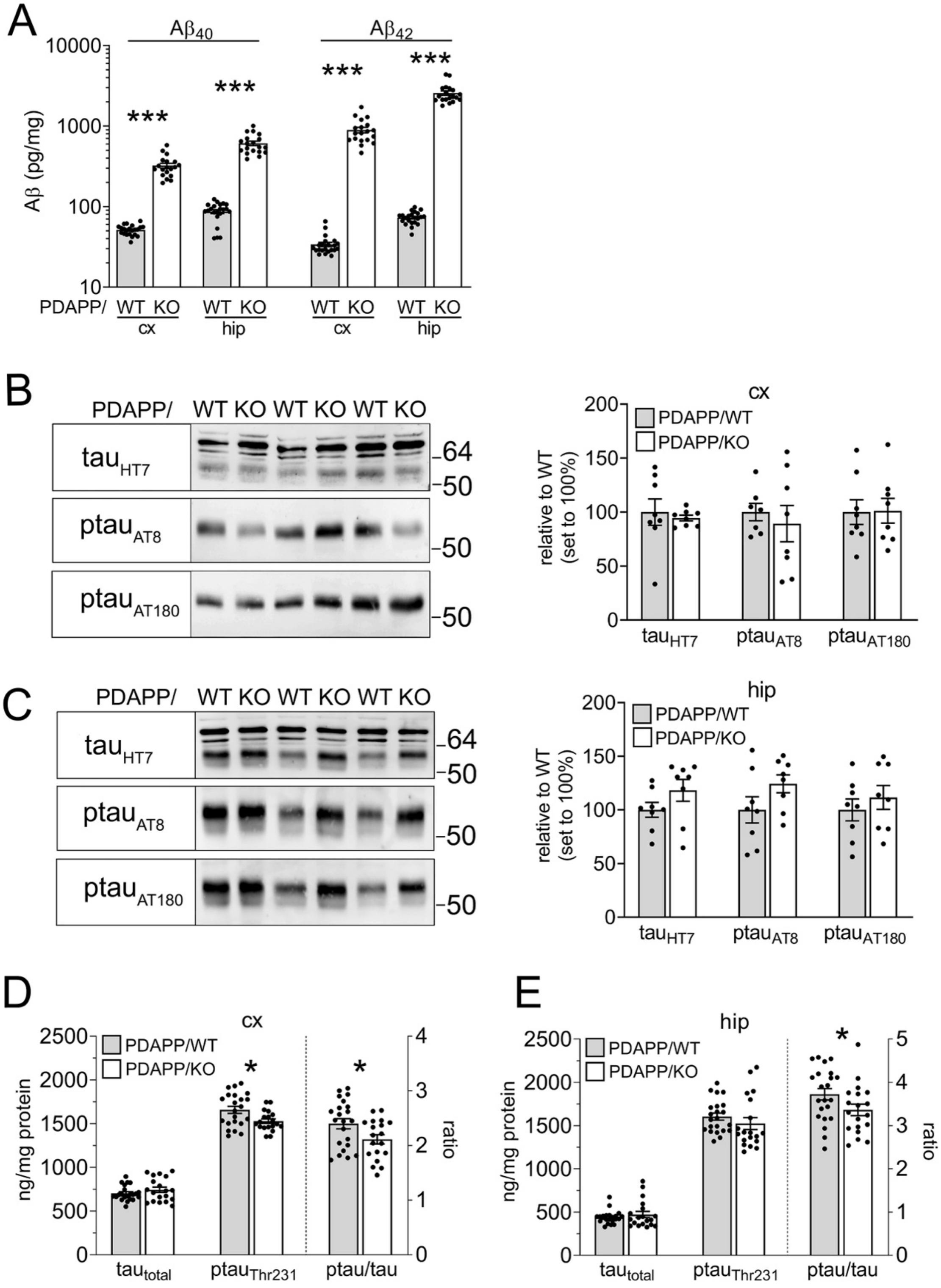


**Figure S2: Amyloid and tau phenotypes in male PDAPP mice lacking SORCS2**

**(A)** Levels of soluble $A\beta_{40}$ and $A\beta_{42}$ in cortex (cx) and hippocampus (hip) of 40 weeks old PDAPP/WT and PDAPP/KO males. Data are given as mean ± SEM from n=19-20 animals per genotype (unpaired Mann-Whitney U-test). **(B, C)** Representative Western blots (left panels), and densitometric quantification of replicate blots (right panels), documenting levels of total tau (HT7) as well as phosphorylated variants $ptau_{Ser202/Thr205}$ (AT8) and $ptau_{Thr231}$ (AT180) in cx (B) and hip (C) of 40 weeks old PDAPP/WT and PDAPP/KO males. Data are expressed as relative to WT (set to 100%) and given as mean ± SEM from n=8 animals per genotype (unpaired Mann-Whitney U-test or two-sided unpaired Student´s *t*-test). **(D, E)** Levels of total ($tau_{total}$) and phosphorylated variants ($ptau_{Thr231}$) of tau, as well as ratio of $ptau_{Thr231}/tau_{total}$, in cx (D) and hip (E) of 40 weeks old PDAPP/WT and PDAPP/KO males as measured by ELISA. Data are given as mean ± SEM from n=19-20 animals per genotype (unpaired Mann-Whitney U test) *, $P < 0.05$; ***, $P < 0.001$

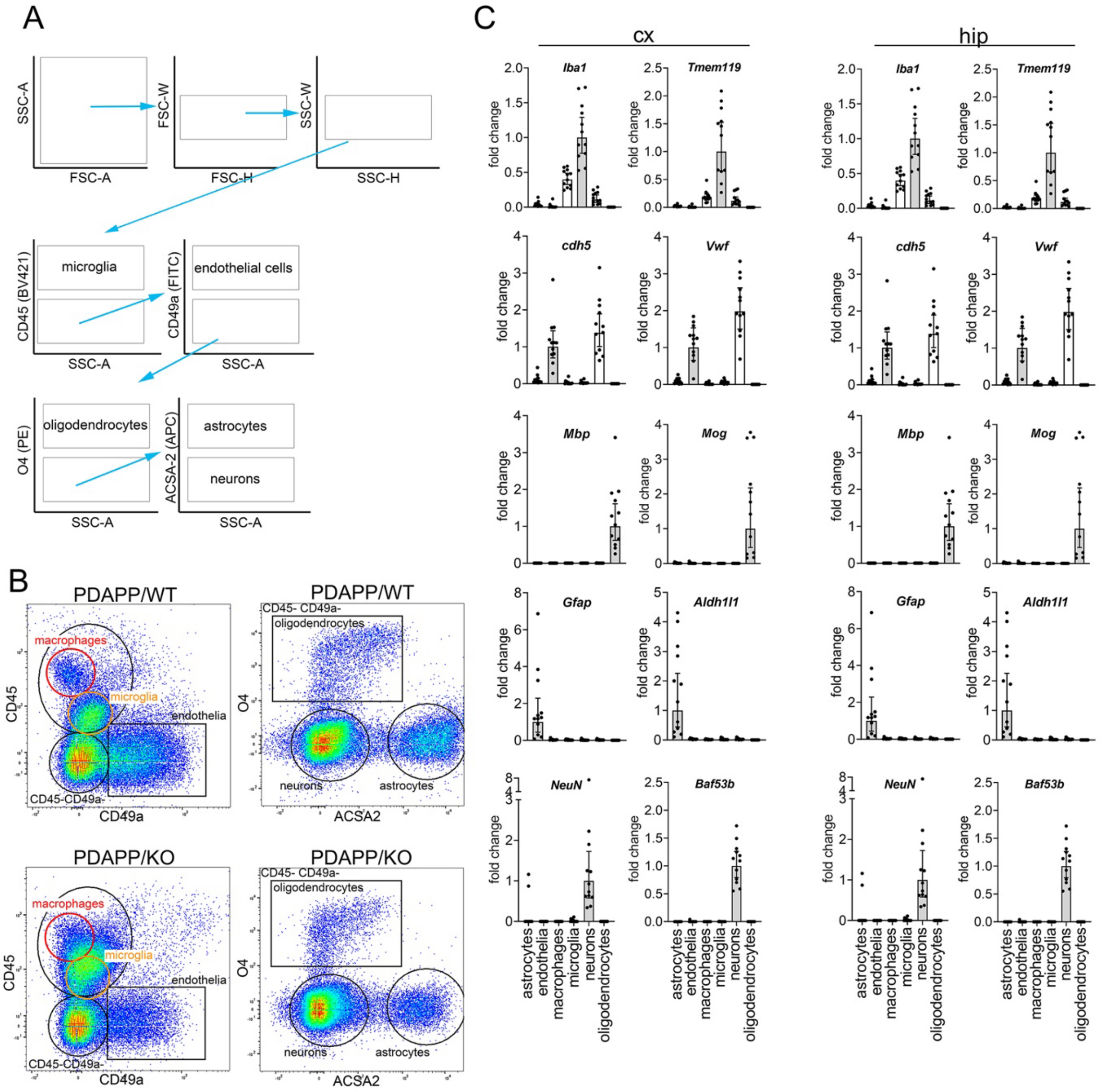


**Figure S3: Protocol for FACS of cell types from adult mouse brains**

**(A)** Strategy for multicolor FACS of cells from adult mouse brains. Isolation of cell types was performed using markers CD45+ (microglia, macrophages), CD45- CD49a+ (endothelia), CD45- CD49a- O4+ (oligodendrocytes), CD45- CD49a- O4- ACSA2+ (astrocytes), and CD45- CD49a- O4- ACSA2- (neurons). **(B)** Representative FACS panels for cortex from PDAPP/WT and PDAPP/KO females using markers CD45 (macrophages, microglia), CD49a (endothelia), O4 (oligodendrocytes), and ACSA2 (astrocytes). The remaining cell population is enriched in neurons. **(C)** The identity of sorted cells from cx and hip samples (pool of PDAPP/WT and PDAPP/KO) was established using qRT-PCR. Markers identify astrocytes (*Gfap, Aldh1l1*), endothelia (*cdh5, Vwf*), macrophages/microglia (*Iba1, Tmem119*), oligodendrocytes (*Mbp, Mog*), and neurons (*NeuN, Baf53b*). *GapdH* was used as internal control. In each cell population, transcript levels of the markers are given as relative to the marker gene symbolized by the grey bar (*e.g., Iba1* in microglia). Data are given as mean ± SEM for n=6 animals per genotype.

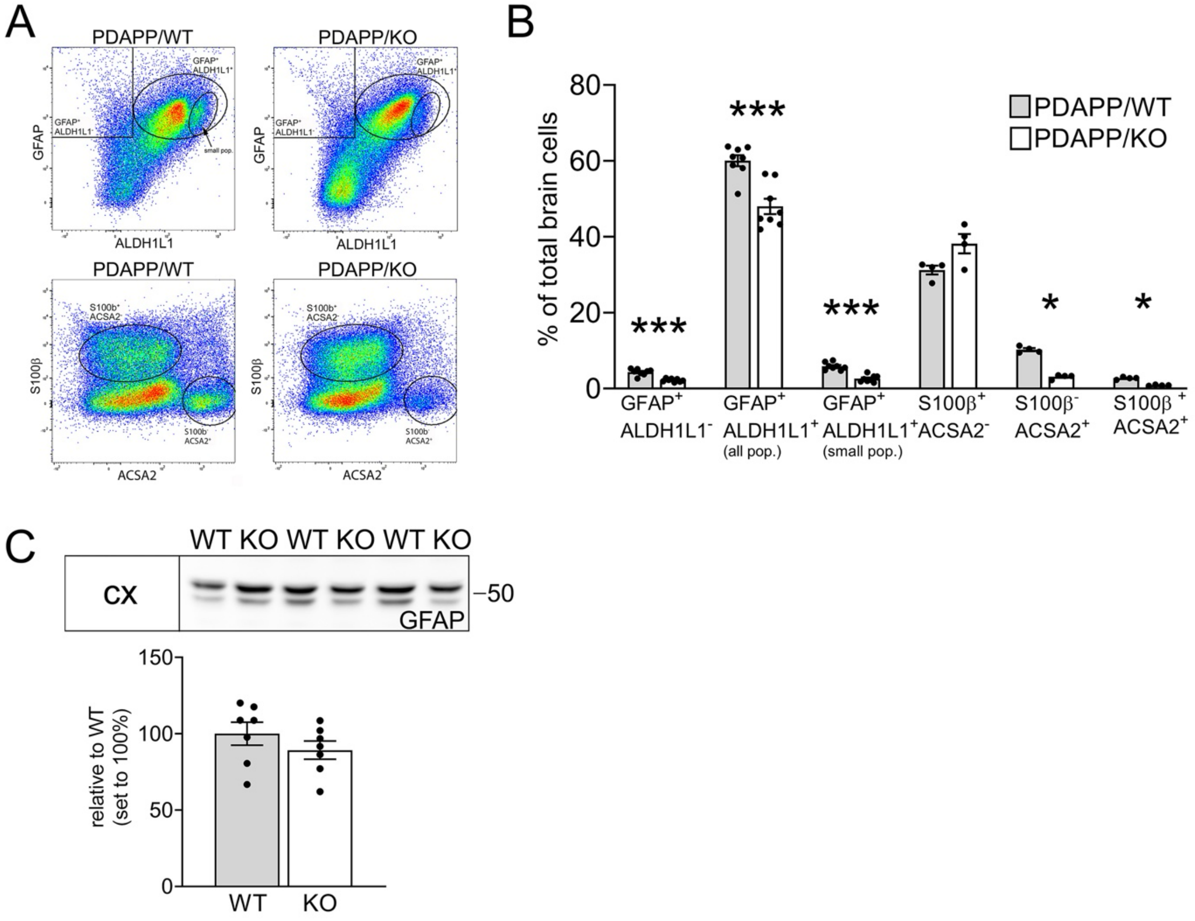


**Figure S4: Aβ induces loss of astrocytes in SORCS2-deficient mice**

**(A, B)** Exemplary dot blots (A), and quantifications thereof (B), showing FACS staining profiles and gating (black boxes or circles) of astrocyte subpopulations in cortices of 40 weeks old PDAPP/WT and PDAPP/KO mice using markers GFAP, ALDH1L1, S100β, and ACSA2. Data in B are given as mean ± SEM for n=4-8 animals per genotype (two-sided unpaired Student´s *t*-test for fractions stained with GFAP, unpaired Mann-Whitney U test for fractions stained with S100β). **(C)** Representative Western blot for GFAP in cx extracts (upper panel), and quantification from densitometric scanning of replicate blots (lower panel), showing comparable levels in 40 weeks old WT and KO mice lacking PDAPP. Data are expressed as relative to WT (set to 100%) and given as mean ± SEM for n=7 animals per genotype (unpaired Mann-Whitney U test). *, $P < 0.05$; ***, $P < 0.001$

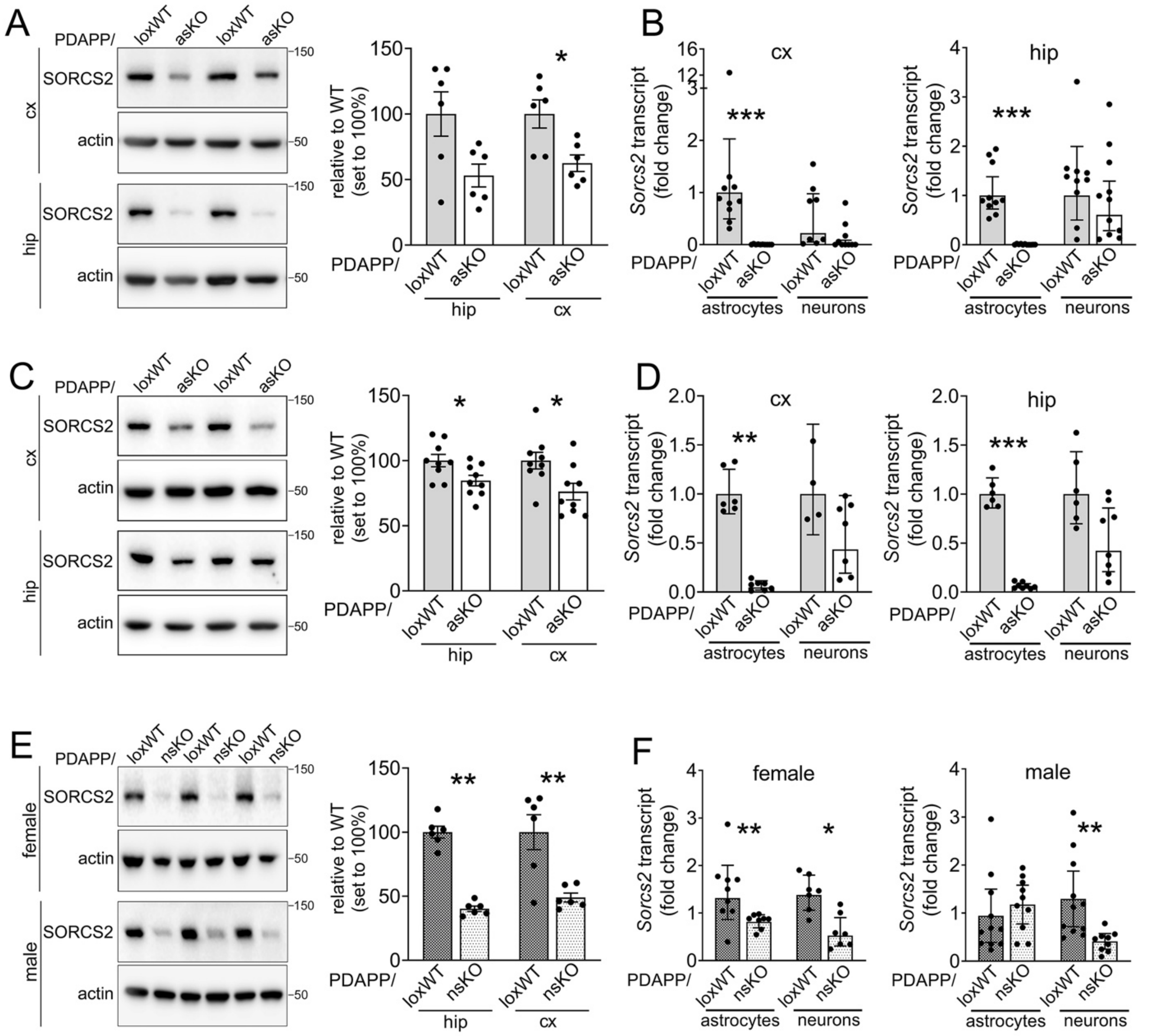


**Figure S5: Validation of mouse models carrying astrocyte- or neuron-specific *Sorcs2* defects**

**(A)** Western blot analyses of cortical (cx) or hippocampal (hip) brain extracts document reduction in SORCS2 levels in female PDAPP mice with astrocyte-specific *Sorcs2* defect (PDAPP/asKO) as compared to PDAPP/loxWT. Animals were 38 weeks of age with time of Cre-ERT2 induction at 26 weeks of age. Exemplary blots as well as quantifications from densitometric scanning of replicate blots are shown. Detection of actin served as loading control. Data are expressed as relative to WT (set to 100%) and are given as mean ± SEM from n=6 animals per genotype (unpaired Mann-Whitney U-test). **(B)** Levels of *Sorcs2* transcript in astrocytes and neurons sorted from cx and hip of PDAPP/loxWT or PDAPP/asKO mice at 38 weeks of age. Loss of *Sorcs2* transcript is seen in astrocytes, but not neurons, in both brain regions. Data are expressed as relative to loxWT (set to 1) and given as mean ± 95% confidence interval from n=6-11 animals per genotype (unpaired Mann-Whitney U-test). **(C, D)** Experiments as in A-B, but testing PDAPP/loxWT and PDAPP/asKO females at 30 weeks of age (induction of Cre-ERT2 at 20 weeks of age). Data are given as mean ± SEM (C) or mean ± 95% confidence interval (D) from n=6-9 animals per genotype (unpaired Mann-Whitney U-test). (**E**) Western blotting and densitometric scanning of replicate blots, show reduction in SORCS2 levels in female and male PDAPP mice with neuron-specific *Sorcs2* defect (PDAPP/nsKO) compared to PDAPP/loxWT animals at 38 weeks of age. Detection of actin served as loading control. Data are expressed as relative to WT (set to 100%) and given as mean ± SEM from n=6 animals per genotype (unpaired Mann-Whitney U-test). **(F)** *Sorcs2* transcript levels in astrocytes and neurons sorted from cx of PDAPP/loxWT or PDAPP/nsKO mice at 38 weeks of age. In PDAPP/nsKO males, *Sorcs2* transcript levels are reduced in astrocytes but not neurons, while being reduced in both cell types in PDAPP/nsKO females. Data are expressed as mean ± 95% confidence interval from n=8-11 animals per genotype (two-sided unpaired Student´s *t*-test). *, P < 0.05; **, P < 0.01; ***, P < 0.001

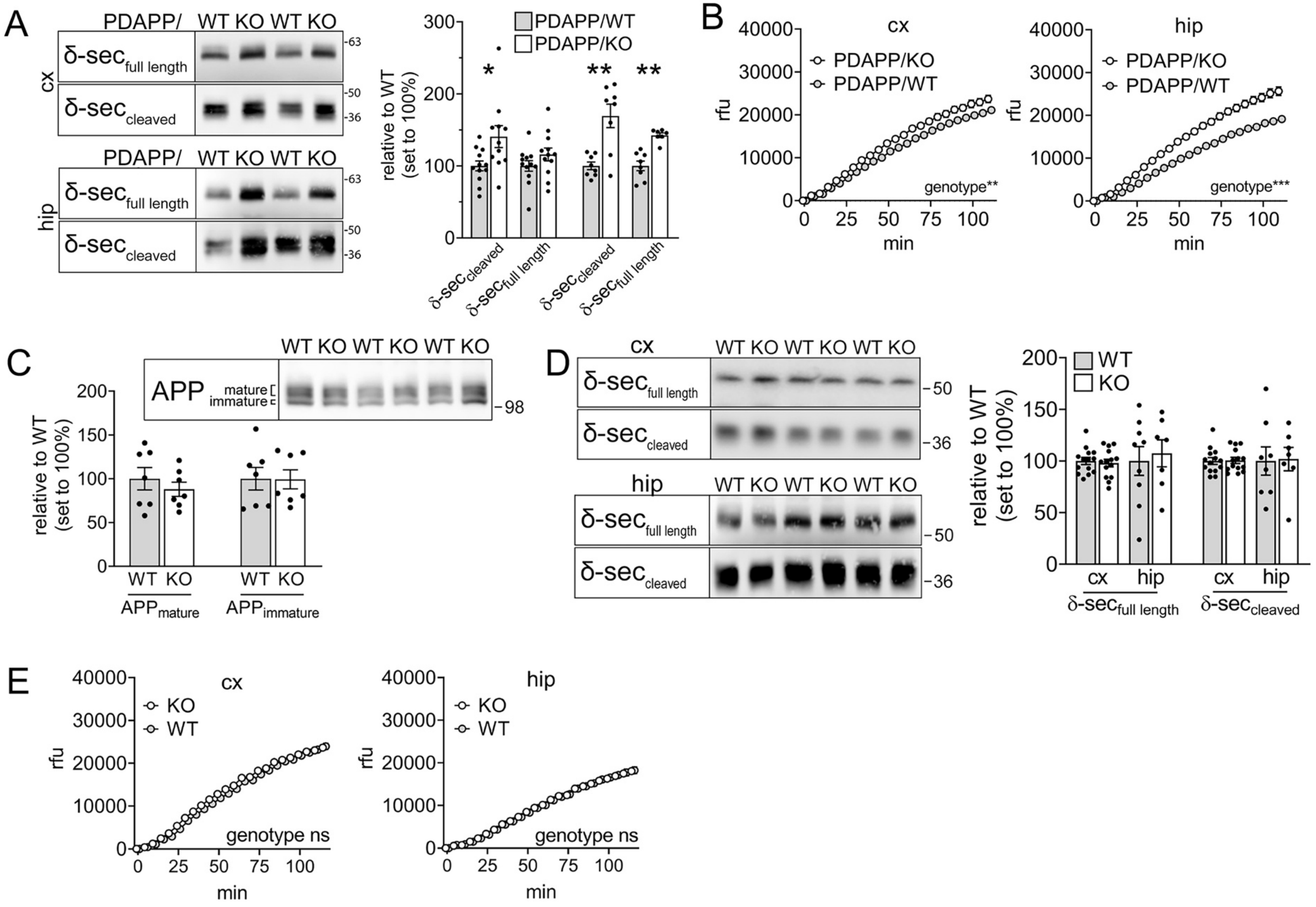


**Figure S6: Induction of δ secretase expression and activity requires Aβ trigger**

**(A)** Representative Western blot analysis, and densitometric scanning of replicate blots thereof, document increased levels of full-length and cleaved forms of δ secretase in cortical (cx) and hippocampal (hip) brain extracts of 40 weeks old PDAPP/KO compared with PDAPP/WT males. Data are expressed as relative to WT (set to 100%) and given as mean ± SEM from n= 7-12 animals per genotype (two-sided unpaired Student´s *t*-test). (**B**) Activity of δ secretase, as determined by fluorogenic substrate cleavage assay, is significantly increased in cx and hip extracts of 40 weeks old PDAPP/KO as compared to PDAPP/WT male mice. Data are given as mean ± SEM from n=8-10 animals per genotype (matched Two-way ANOVA). (**C**) Representative Western blot and densitometric scanning of replicate blots, show comparable levels of immature and mature APP in cx extracts of 40 weeks old WT and KO females lacking PDAPP. Data are given as mean ± SEM from n=7 animals per genotype (unpaired Mann-Whitney U-test). (**D**) Experiment as in A, showing comparable levels of full length and cleaved forms of δ secretase in cx and hip extracts of 40 weeks old WT and KO females lacking PDAPP. Data are given as mean ± SEM from n= 7-14 animals per genotype (unpaired Mann-Whitney U-test for hip, two-sided unpaired Student´s *t*-test for cx). (**E**) Experiment as in B, documenting comparable δ secretase activity in cx and hip extracts from 12 weeks old PDAPP/WT and PDAPP/KO females. Data are given as mean ± SEM from n=6 animals per genotype (matched Two-way ANOVA). ns. not significant; rfu, relative fluorescence units. *, $P < 0.05$; **, $P < 0.01$

### SUPPLEMENTARY TABLE

**Table S1:** Inflammatory profiles of cortical lysates from 20 and 40 weeks old PDAPP/WT and PDAPP/KO females. Aβ levels determined for all animals are given as concentration range for the respective cohort. Young females were grouped into two cohorts, having modest or high levels of Aβ (cut off: 300 pg/ml $A\beta_{40}$ and 800 pg/ml $A\beta_{42}$ for hip; 35 pg/ml $A\beta_{40}$ and 60 pg/ml $A\beta_{42}$ for cx). Data are given as mean ± SD from n=11-22 animals per genotype (two-sided unpaired Student´s *t*-test). ns, not significant; N/A, not available. Markers significantly increased or decreased in PDAPP/KO as compared to matched PDAPP/WT females are highlighted in red or green, respectively.

| | 20 weeks | | | 40 weeks | | 20 weeks | | 40 weeks |
|---|---|---|---|---|---|---|---|---|
| **Analyte** | **WT** | **KO (low Aβ)** | **KO (high Aβ)** | **WT** | **KO** | **P value** | | |
| | **pg/mg** | **pg/mg** | **pg/mg** | **pg/mg** | **pg/mg** | **WT vs KO (low Aβ)** | **WT vs KO (high Aβ)** | **WT vs KO** |
| **pro-inflammatory cytokines / chemokines** | | | | | | | | |
| IL-1β | 1.43±0.60 | 1.84±0.44 | 0.94±0.41 | 7.49±4.01 | 3.60±1.42 | 0.05 | <0.05 | <0.0001 |
| TNFα | 1.12±0.24 | 1.58±0.29 | 1.08±0.20 | 5.70±4.31 | 5.78±2.01 | <0.001 | ns | ns |
| IL-12p70 | 69.09±21.74 | 103.20±26.13 | 48.71±12.82 | 103.90±58.87 | 47.60±15.99 | <0.01 | <0.01 | <0.001 |
| IL-6 | 48.01±18.84 | 68.72±14.34 | 31.52±15.22 | 15.47±7.80 | 7.51±2.43 | <0.01 | <0.05 | <0.0001 |
| IP10/CXCL10 | 3.54±0.47 | 6.14±2.30 | 7.80±2.05 | 21.99±14.62 | 41.38±12.88 | <0.01 | <0.0001 | <0.0001 |
| MIP1α/CCL3 | 1.64±0.23 | 4.33±1.65 | 5.27±1.16 | 8.42±6.22 | 20.76±5.68 | <0.001 | <0.0001 | <0.0001 |
| IL-2 | 1.94±0.82 | 2.78±0.56 | 1.57±0.39 | 5.39±3.54 | 2.56±0.84 | <0.01 | ns | <0.01 |
| IL-5 | 2.85±0.97 | 3.97±1.19 | 1.98±0.88 | 0.77±0.43 | 0.37±0.13 | <0.05 | <0.05 | <0.001 |
| IL-16 | 192.00±29.41 | 209.70±41.11 | 277.90±112.0 | 108.30±82.06 | 538.3±155.09 | ns | <0.05 | <0.0001 |
| YKL40/CHI3L1 | 0.63±0.13 | 0.93±0.28 | 1.37±0.36 | 3.393±2.25 | 10.47±5.29 | <0.0001 | <0.0001 | <0.0001 |
| **anti-inflammatory cytokines** | | | | | | | | |
| IL-10 | 2.62±0.85 | 3.03±0.44 | 1.95±0.84 | 7.22±3.40 | 2.71±1.29 | ns | <0.05 | <0.0001 |
| IL-4 | 1.02±0.35 | 1.22±0.41 | 0.65±0.28 | 1.02±0.45 | 0.38±0.18 | ns | <0.01 | <0.0001 |
| IL-22 | 6.87±0.87 | 6.73±0.65 | 4.58±0.89 | 10.48±6.03 | 5.58±2.18 | ns | <0.0001 | <0.01 |
| TGFβ2 | N/A | N/A | N/A | 12.24±0.45 | 9.93±2.11 | N/A | N/A | <0.001 |
| TGFβ3 | N/A | N/A | N/A | 1.15±0.21 | 0.75±0.15 | N/A | N/A | <0.0001 |
